# Multiplexed single-cell proteomics using SCoPE2

**DOI:** 10.1101/2021.03.12.435034

**Authors:** Aleksandra A. Petelski, Edward Emmott, Andrew Leduc, R. Gray Huffman, Harrison Specht, David H. Perlman, Nikolai Slavov

## Abstract

Many biological systems are composed of diverse single cells. This diversity necessitates functional and molecular single-cell analysis. Single-cell protein analysis has long relied on affinity reagents, but emerging mass-spectrometry methods (either label-free or multiplexed) have enabled quantifying over 1,000 proteins per cell while simultaneously increasing the specificity of protein quantification. Isobaric carrier based multiplexed single-cell proteomics is a scalable, reliable, and cost-effective method that can be fully automated and implemented on widely available equipment. It uses inexpensive reagents and is applicable to any sample that can be processed to a single-cell suspension. Here we describe an automated Single Cell ProtEomics (SCoPE2) workflow that allows analyzing about 200 single cells per 24 hours using only standard commercial equipment. We emphasize experimental steps and benchmarks required for achieving quantitative protein analysis.

**SCoPE2 Protocol:** 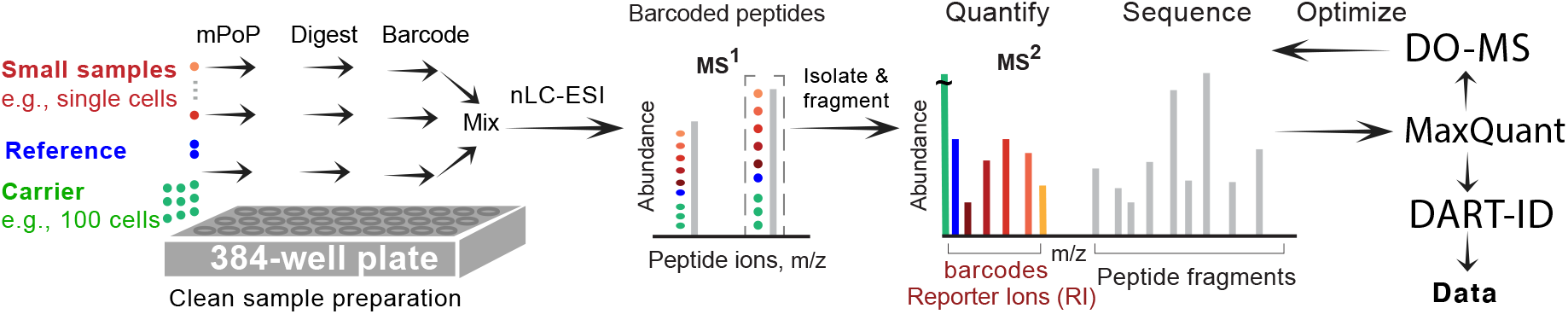

## Introduction

Biological systems, such as the tissues of multicellular organisms, are composed of diverse cell types and states^1–4^. This cellular diversity is well appreciated and has motivated the fruitful development of numerous analytical approaches for analyzing individual cells^2–5^. Indeed over the last decade, multiplexed approaches for single-cell transcriptomics have scaled to detecting the transcripts from thousands of genes across many thousands of single cells^5^. Single-cell transcriptomics methods are proving useful in understanding fundamental and clinical problems, such as the interaction of cancer and immune cells^6^ and cancer drug resistance^7^. Despite this progress, the pervasive post-transcriptional regulation across human tissues^8^ cannot be characterized based on nucleic acids analysis alone and has motivated methods for analysing proteins in single cells^9^.

Traditionally, single-cell protein analysis has relied primarily on affinity reagents, such as antibodies and aptamers^3^ while the powerful mass-spectrometry (MS) methods that afford comprehensive quantification of cellular proteomes have been limited to the analysis of bulk samples composed of many cells^10–14^. Recently, new MS methods have been developed for quantifying thousands of protein in individual human cells as reviewed in refs.^15,16^. These single-cell mass-spectrometry methods hold much potential to facilitate the characterization of molecular mechanisms of health and disease^9^.

### Development of Single Cell ProtEomics (SCoPE2)

SCoPE2 is a second generation method enabled by concepts and approaches introduced by its first generation method (Single Cell ProtEomics by Mass Spectrometry; SCoPE-MS^17^) and new ones introduced with SCoPE2^18^. For example the isobaric carrier concept^19^ was first developed and introduced for multiplexed single-cell analysis as part of SCoPE-MS while data analytics (described below) for optimizing experimental designs and enhancing data interpretation were introduced with SCoPE2^18^. Some concepts shared by SCoPE-MS and SCoPE2 (such as the use of a clean cell lysis that obviates clean-up and associated sample losses) have different implementations: SCoPE-MS implemented this idea via adaptive focused acoustics^17^ while SCoPE2 implemented it as Minimal ProteOmic sample Preparation (mPOP)^20^. mPOP allows to automate sample preparation while reducing its volumes. Recently, we introduced a droplet method, nano-ProteOmic sample Preparation (nPOP), that enabled an automated and simultaneous preparation of hundreds of single cells in 20 nl droplets^21^. As a whole, the second generation method, SCoPE2, has increased throughput and quantitative accuracy while lowering the cost and the barriers to adoption^15,18^.

The isobaric carrier concept is central to SCoPE2. It involves labeling peptides from individual cells with isobaric mass tags and combining them with ‘isobaric carrier’ peptides derived from a larger number of cells to reduce surface adsorption losses of single-cell peptides and to provide peptide fragments enhancing peptide sequence identification^19,22^. Importantly, the isobaric carrier approach allows designing experiments to maximize either depth of proteome coverage or to maximize copies of ions sampled per peptide^19^. The isobaric carrier approach has been adopted by multiple laboratories for ultra-sensitive MS analysis of single cells and other small samples (reviewed in ref.^15^).

To enable inexpensive and robust single-cell proteomics, SCoPE2 built upon key ideas of SCoPE-MS and introduced many technological and analytical improvements^18^. Similar to SCoPE-MS, SCoPE2 uses clean cell lysis in water, but replaced the low-throughput focused acoustic sonication with mPOP, a lower volume and higher throughput freeze-heat cycle that enabled lysing cells in multi-well plate formats^18,20,23^. We also enhanced the means of normalising single-cell quantification data by the inclusion of a 5-cell reference channel, prepared in bulk and common across all SCoPE2 sets from an experiment^18^. Some SCoPE2 spectra allow quantifying the corresponding peptides but do not contain enough peptide fragments to support confident sequence identification. To help recover these additional peptides, we introduced the DART-ID software available from dart-id.slavovlab.net. It implements a false discovery rate controlled Bayesian update of peptide confidence of identification by employing informative features of peptides, for example its aligned retention time across many LC-MS/MS runs^24^. To optimize instrumentation settings for SCoPE2 sample analysis, we developed an approach to Data-driven Optimization of MS (DO-MS)^25^. This approach is implemented by extendable Shiny interface freely-available from do-ms.slavovlab.net. DO-MS allows analyzing and optimising LC-MS/MS experiments, with particular utility for ultra-sensitive proteomics as employed in SCoPE2^18,25^.

In this protocol, we provide a detailed description of the principles and experimental steps that allow adopting and applying SCoPE2 to different systems. We have already published software tools^24,25^ and experimental guidelines^19^ to aid the adoption of SCoPE2. However, fulfilling its potential for making quantitative measurements requires a comprehensive guide to designing, implementing, and benchmarking single-cell mass-spec analysis. Here, we aim to provide such a guide, emphasizing the principles that enable quantitative SCoPE2 analysis and tailoring it to different experimental constraints and scientific aims.

### Applications of SCoPE2

Any samples of primary tissues or cell cultures that can be prepared as a suspension of single cells can be analyzed by the SCoPE2 protocol described here. The single-cell suspensions can be prepared by methods used for single-cell RNA-seq. While most methods should be as applicable to preparing single cells for SCoPE2 analysis, some methods, such as protease treatments, might affect cell surface proteins. Furthermore, the SCoPE2 protocol described here and its principles for ultra-sensitive analysis may be applied successfully to other small samples, such as biopsies.

### Comparison with other methods for single-cell protein analysis

Single-cell methods for investigating protein levels have existed for decades in the form of technologies employing affinity reagents (such as antibodies) and fluorescent proteins [3]. These classical approaches either employ antibodies against epitopes of interest, or use modified cells expressing fluorescent fusion proteins or reporters for a protein of interest. These methods therefore require antibodies which vary in their specificity, or engineering fluorescent fusion proteins or reporters which may influence the activity of a protein of interest or its modified host cell [3]. Both approaches are limited in the number of proteins that can be analyzed to about 1-10 target proteins. This limit has been relaxed to about 50-100 target proteins by advanced methods barcoding the affinity reagents. An example of such barcoding include CyTOF which uses antibodies conjugated to rare-earth metals^26^. Other examples include approaches such as REAP-seq and CITE-seq that use DNA olgo-linked antibodies, and permit higher multiplexing, as well as the possibility of obtaining both single-cell level RNA and protein data^3^. The limited specificity of affinity reagents can be mitigated by targeting multiple epitopes per proteins, as implemented by proximity extension and ligation assays^27,28^, or by using additional features of the protein, as implemented by single-cell western blots^29^. These advances have enabled approaches based on affinity reagents to quantify about a 100 proteins per single cell. However highly specific antibodies and antibody validation remain required and sometimes challenging steps for these approaches.

Mass-spectrometry-based approaches offer an alternative, bringing the promise of quantifying orders of magnitude more proteins, without the time and expense of obtaining and qualifying specific antibodies, or the potential to inadvertently disrupt normal protein function through generation of fusion fluorescent proteins^9^. Two main types of MS methods have been introduced:(i) multiplexed approaches employing isobaric carriers, including SCoPE-MS^17^ and SCoPE2^18,30^ discussed here, and label-free approaches^31–35^. The label-free approaches often seek to miniaturise traditional proteomic sample preparation to enable processing of individual single cells with minimal sample losses due to the sub-microlitre sample preparation volumes used by approaches such as nanoPOTs^36,37^ and OAD^38^. Recently, some of these approaches have been automated^39^; for comprehensive reviews, see refs.^15,16^. Each single-cell lysate is then digested and analysed individually by LC-MS/MS, usually using MS1-based quantification. In contrast, the SCoPE2 protocol presented here sought to minimize losses during chromatography by using an isobaric carrier and to avoid the losses inherent in sample cleanup procedures through the use of clean lysis. Importantly, the SCoPE2 approach uses sample multiplexing, so that protein identification can be performed on material pooled from multiple cells, rather than a single-cell, with TMT 16-plex reagents permitting multiplexed analysis of 12-14 single cells per experiment. Detailed reviews of current MS methods to conduct single-cell proteomics can be found in refs.^15,16^.

### Experimental design

Design considerations familiar to scRNAseq practitioners are also applicable to SCoPE2 experiments^40^. Those parameters in common include the scale of the experiment and distributing populations of interest across batches to minimise the impact of batch effects. Similarly many of the downstream types of data analysis steps, such as batch correction, imputation and dimensionality reduction, are similar to those used for processing scRNAseq data. Indeed, packages developed for performing procedures such as multi-single-cell ‘omics dataset alignment and pseudotime or trajectory inference are compatible with single-cell proteomics. For example, we used Conos^41^ for joint analysis of RNA and proteins^18^. We needed to implement slight adaptations to Conos to import SCoPE2 data. Similarly, other methods might need minor adjustments. Furthermore, methods developed for scRNA-seq do not take advantage of specific features to single-cell proteomics data^15,42^. These design considerations are common to many different forms of single-cell omics analysis and are discussed extensively elsewhere^40,43^. Several parameters should be considered that are specific to, or should be considered differently for SCoPE2-based single-cell proteomics analysis:

- **Single-cell isolation:** Similarly to scRNAseq, it is necessary to disaggregate complex samples or tissues prior to SCoPE2 analysis in order to obtain a single-cell suspension which can then be further processed, for example by FACS or CellenONE. While enzymatic treatment such as the use of trypsin or accutase are common, these can influence the surface proteome, so the method of choice may differ between scRNAseq and SCoPE2 sample preparation depending on the sample and experimental question.
- **Single-cell population selection:** Enrichment of rare, or specific subpopulations of interest is often performed for scRNAseq, though is especially pertinent for SCoPE2 experiments. It is challenging for SCoPE2 experiments to reach the very high numbers of cells that some of the current scRNAseq methods can yield. Enriching particular subpopulations of interest is a useful strategy to help mitigate this issue, but will influence the selection of carrier and reference proteome composition.
- **Carrier and reference composition:** Single-cell sampling for SCoPE2 experiments can allow for enrichment of specific subpopulations of interest. However, when performing SCoPE2 experiments in a data-dependent manner, the carrier and reference channels should be an even mix of the populations of interest or a close to it as is achievable. Uneven representation in the carrier or reference could result in failure to detect proteins only represented in particular subpopulations. One potential mechanism to alleviate this issue is to spike specific peptides of interest into both carrier and reference channels. Such spike-ins will ensure that those peptides are sent for MS2 analysis. For example, using carrier peptides enriched for post-translational modifications (PTMs, e.g., phosphorylation) or adding synthetic peptides may enable single-cell PTM analysis as previously suggested^9^. However, if the abundance of these peptides in the single cells is very low, they may not be quantified in the single cells. This challenge may be partially mitigated by increasing the MS2 accumulation time in order to increase the chances of quantifying those peptides in the single cells. While spiking in synthetic peptides also adds cost to performing SCoPE2 experiments, it may be well justified by the increased probability of analyzing proteins and modifications of special biological interest.
- **Carrier abundance:** While a 100-cell carrier has proven suitable for a majority of our experiments, this is a parameter that will likely require adjusting depending on the cells of interest and their proteome abundance. Optimization of carrier abundance is discussed in the protocol. Higher carrier abundance can allow identifying peptides with shorter ion accumulation times, and thus analyzing more peptides per unit time. However, the shorter accumulation times will reduce the ion copies sampled from single-cell proteins and thus will reduce quantitative accuracy and increase missing data in single cells. An extensive discussion of considerations for carrier abundance and optimisation for different experimental designs can be found in Specht & Slavov (2020)^19^.
- **Single or dual carrier selection:** The use of a single carrier channel maximizes the number of single cells that can be analysed per SCoPE2 run. Depending on the experimental design, however, the use of a second carrier channel can provide a valuable internal control, and we describe such an example here with the 100xMaster samples for LC-MS/MS optimization.
- **Channel Selection for Carrier:** The isobaric tag used to label the carrier peptides should be chosen to minimize its influence on samples labeled with other tags. This includes two considerations. First, the isotopic contaminations of the tag should affect as few samples as possible, which is the case for the lightest and the heaviest tags since their isotopic contaminants affect only one sample. Second, ideally the tag should not be mDa away from another tag (e.g., a C-N pair tag) since these tags become challenging to resolve when used for samples of very different abundance. These considerations favor the use of 126 with TMT 11-plex (since 131 is a C-N pair) and 126 or 134N with TMTpro 16-plex since 134C is not used for the 16-plex.
- **LC-MS/MS system suitability:** Before conducting a SCoPE2 experiment, an LC-MS/MS system can require significant optimisation to allow it to successfully obtain meaningful SCoPE2 data. We recommend the use of dilute 100xMaster standards, diluted to single-cell levels to optimise instrument performance without the biological variation inherent to true single-cell samples. Details on 100xMaster generation and guidelines for instrument optimisation are provided in the SCoPE2 protocol below.
- **Positive and Negative Controls within single cell sets:** We recommend adding controls within each SCoPE2 experiment. Positive controls allow to evaluate sample sample preparation independent of cell isolation and are particularly useful when the quality of the single cells or their isolation are uncertain. Negative controls allow to evaluate background noise and are particularly useful for evaluating potential problems with sample preparation, such as cross labeling.

### Level of expertise needed to implement the protocol

The sample handling portions of this protocol should be approachable for a biochemist or cell biologist with cell culture and molecular biology experience. It is recommended that users not experienced in sample preparation for proteomics consult with an individual with experience in this area to avoid common mistakes such as polymer contamination deriving from plasticware or improperly handled buffers.

The LC-MS/MS system requires the engagement of an experienced mass spectrometrist early in the project to ensure that the instrumentation is performing at a level suitable for single-cell analysis. Many core facilities will not be able to perform this analysis without extensive planning and engagement. Due to the range of samples processed, facility workflows are typically optimised for robustness as opposed to sensitivity. Due to the extremely low abundance of single cell samples and carryover from more abundant samples run previously on the LC-MS/MS system, significant optimisation may be required to adapt workflows, though guidelines for this process are described in the protocol.

Data analysis steps specific to SCoPE2 single-cell proteomics experiments are presented, for example quality control metrics and identification of failed wells. Downstream analysis including batch correction, dimensionality reduction, and differential expression falls outside the scope of this protocol, but will be familiar to users with experience in scRNAseq data processing.

## Limitations

The current sample preparation method described here is robust and uses equipment readily available to many labs. However, while the volumes used for sample preparation represent an order of magnitude decrease from the original SCoPE-MS protocol^17^, they are still orders of magnitude larger than those used for droplet based scRNAseq sample processing. Reduced sample volumes will significantly reduce sample losses to surfaces during processing. Alternative approaches for preparing single cells for mass-spec analysis, such as OAD, and nanoPOTs, allow for reducing sample processing volumes to 200 nL^36^, a 5 to 10-fold reduction on the 1-2 *µL* used here. If desired, OAD and nanoPOTs can be easily incorporated within the SCoPE2 framework. However, these methods have not been scaled efficiently to hundreds of cells the way mPOP has been.

Currently a major bottleneck for all single-cell mass-spectrometry methods is the large number of LC-MS/MS runs required to analyze thousands of single cells. The multiplexing afforded by SCoPE2 reduced the number of runs needed by over 10 fold, but the number remains large and likely represents the single largest expense for the analysis, particularly for investigators working through a core facility^18^. Improved barcoding strategies, for example the newly announced TMT-pro reagents from Thermo Scientific allows at least 50% increase in throughput by reducing the number of required LC-MS/MS runs by approximately 1/3 for analysis of the same number of single cells. A strategy that can further increase multiplexing is the use of both TMT 11plex and TMTpro 16plex reagents. This strategy offers trade offs discussed below. The possibilities and limitations of increasing multiplexing further are discussed in ref.^15^.

Coisolation and co-fragmentation of ions limits the quantitative accuracy of isobaric labeling methods, including SCoPE2. This limitation can be mitigated by performing quantification at the MS3 level at the cost of sensitivity loss. Another solution would be to use complement ions, which are peptide fragments with the balancer portion of the TMT still attached. In our laboratory, we aim to mitigate coisolation by sampling peptides at the apex of the elution peaks to reduce coisolation, along with using narrow isolation windows. The optimization of the apex targeting is facilitated by DO-MS and has the additional benefit of increasing the number of ions copies samples per unit time and thus sensitivity and quantification accuracy^25^.

Another limitation originates from the tendency of current workflows to sample only a small fraction of ions available for analysis. Such sampling has been highly successful in bulk proteomics where only short accumulation times are typically required to sample individual peptides. The ability to sample ions more completely, e.g., by accumulating ions in parallel and thus for longer times (as exemplified by the PASEF mode of the timsTOF instruments^44^) can substantially enhance the sensitivity and the accuracy of quantification. A trade-off of the timsTOF instruments at present is that their resolution does not permit the level of sample multiplexing possible with Orbitrap instruments and TMT-based isobaric labelling reagents, thus limiting the number of cells that can be analysed per LC/MS-MS run.

These limitations and how they may be mitigated by future developments are discussed in more details in ref.^45^. These developments will further improve the sensitivity, throughput and robustness of single-cell protein analysis.

## Materials

### Biological Materials

- Isolated primary cells, or cell lines grown in culture. **Caution: Cell lines should be checked to ensure they are authentic, and free of mycoplasma contamination**. **Note: While multiple cell lines can be used for preparation of initial SCoPE2 100xMaster samples, our 100x Master samples use U937 (ATCC CRL-1593.2, RRID: CVCL 0007) and Jurkat (ATCC**, **RRID: CVCL 0367) cells, grown in RPMI-1640 medium, supplemented with 10% bovine serum and penicillin/streptomycin**. In order to to generate a small-scale SCoPE2 experiment that is featured within this manuscript, we used U937 and HeLa cells (RRID CVCL 0030).

### Reagents

**Critical: All solutions should be prepared with LC-MS/MS-grade water and reagents. Use of lower quality reagents can result in contamination of solutions with polymers and compromise LC-MS/MS detection of peptides of interest. Plasticware used for storing solvents should be rinsed with ethanol or isopropanol prior to use, and solutions should be used within a week**.

- Water, Optima LC-MS/MS grade (Fisher Scientific; cat. no: W6-1).
- Acetonitrile (for buffer preparation), Optima LC-MS/MS grade (Fisher Scientific; cat. no: A955-1). **Caution: Acetonitrile is a flammable liquid that can irritate the eyes, skin, respiratory tract, central nervous system, and can cause liver or kidney injuries. Wear personal protective equipment when using acetonitrile. This chemical should be handled under a chemical fume hood**.
- Acetonitrile (for Tandem Mass Tag preparation), Anhydrous, 99.8% (Sigma Aldrich; cat. no: 271004-100ML).
- Triethylammonium bicarbonate (TEAB), 1 M pH 8.5 (Sigma Alrich; cat. no: T7408-100ML).
- Formic Acid, Pierce, LC-MS/MS grade (Thermo Fisher Scientific; cat. no: 85178). **Caution: Formic acid is a flammable liquid that can cause serious eye damage or skin burns. Wear personal protective equipment, keep away from any heat, and use in a well-ventilated area**.
- Tandem Mass Tags, TMTpro 16plex Label Reagent Set, 1 x 5 mg (Thermo Fisher Scientific; cat. no: A44520). **Note: we recommend using the 16plex reagents due to their higher throughput, however if you wish to use 11plex reagents they are: Tandem Mass Tags, TMT10plex Isobaric label reagent set plus TMT11-131C (Thermo Fisher Scientific; cat. no: A34808)**.
- Hydroxylamine, 50% w/v (Sigma; 467804-50ML). **Caution: Hydroxylamine can case skin irritation; use appropriate protective equipment**.
- Trypsin, Trypsin Gold Mass Spectrometry Grade (Promega; cat. no: V5280). **Caution: Different sources of trypsin can vary in their purity. Less pure trypsin will negatively impact SCoPE2 results. Caution: Trypsin is chemical that can cause skin, respiratory, and eye irritation. Use under a chemical fume hood with personal protective equipment**.
- Benzonase nuclease, (Sigma Aldrich; cat. no: E1014-25KU).
- MassPREP peptide mixture, (Waters; cat. no: 186002337). **Note: The MassPREP peptide mixture represents a simple mixture of 9 non-tryptic peptides used for passivation of plasticware. It could be substituted with a range of retention time standards or similar mixtures**.
- PBS - Phosphate-Buffered Saline (10X) pH 7.4, RNase-free (Thermo Fisher Scientific; cat. no: AM9625).

### Equipment

- PCR Plate, 384-well, standard (Thermo Fisher Scientific; cat. no: AB1384). **Critical: Different sources of PCR plate can have high levels of polymer contamination rendering them unsuitable for SCoPE2 sample preparation. If using plates from another source, the levels of polymer contamination should be assessed prior to beginning the experiment**.
- Adhesive PCR Plate Foils (Thermo Fisher Scientific; cat. no: AB0626).
- PCR tubes: TempAssure 0.2mL PCR 8-Tube Strips (USA Scientific; cat. no: 1402-3900). **Critical: as described for 384-well plates, different plasticware sources can have high levels of polymer contamination, and this should be assessed before using different PCR strips**.
- Glass autosampler inserts, 9mm (Thermo Fisher Scientific; cat. no: C4010-630). **Critical: The use of glass rather than plastic for sample storage greatly reduces sample losses. The use of autosampler vial inserts permits the SCoPE2 samples to be resuspended and injected into the instrument in smaller (1***µ***L) volumes, again minimising losses**.
- 9mm Clear Glass Screw Thread Vials (Thermo Fisher Scientific; cat. no: 60180-509).
- 9mm Autosampler Vial Screw Thread Caps (Thermo Fisher Scientific; cat. no: C5000-51B).
- 384-well PCR machine with heated lid, e.g. C1000 Touch with 384-well module (Bio-rad; cat. no: 1851138).
- 96-well PCR machine with heated lid, e.g. T100 Thermal Cycler (Bio-rad; cat. no: 1861096), if using this model the use of the tube support ring (Bio-rad; cat. no: 1862000) is recommended). **Note: if only a 384-well PCR machine is available, all steps requiring PCR tubes and a 96-well PCR machine can be accomplished using the 384-well plates in the 384-well PCR machine**.
- Water bath sonicator, e.g. 2.8 L ultrasonic cleaner with digital timer (VWR; cat. no: 97043-964
- Plate Spinner, e.g. PlateFuge microcentrifuge (Benchmark Scientific; Model C2000). **Note: this plate spinner does not offer speed control as it is used to collect liquid at the bottom of a well, rather than for pelleting material**.
- PCR tube spinner, e.g. 16-place microcentrifuge for 0.2mL tubes (USA Scientific; cat. no: 2621-0016).
- Autosampler Vial Spinner, e.g. myFuge 5 (MTC Bio; cat. no: C2595). **Note: this centrifuge does not offer speed control as it is used to collect liquid at the bottom of the autosampler vial, rather than for pelleting material**.
- Vortex, e.g. Analog vortex mixer, (VWR; Model 58816-121).
- Mantis Microfluidic Liquid Handler (Formulatrix) **Caution: If using a different liquid dispensing robot/handler, it is important to check if it is compatible with the 100% acetonitrile that the TMT reagents are in, and also that the plasticware used does not introduce polymer contamination into the samples**.
- Mantis microfluidic chips, low-volume silicone chips (Formulatrix; cat. no: MCLVS12) **Note: These chips are suitable for use with aqueous solutions, and are used for all dispensing steps aside from TMT reagent dispensing as they are not recommended for high solvent concentrations**.
- Mantis microfluidic chips, low-volume 3PFE chips (Formulatrix; cat. no: MCLVPR2) **Note: These chips are suitable for high solvent concentrations and are solely used for dispensing TMT reagents during SCoPE2 sample preparation**.
- Mantis PCR Plate Adapter with wide conical pins for automated plate handling (Formulatrix; cat. no: 232400)
- LC-MS/MS System (e.g. Q-Exactive with Nanospray Flex Ion Source, Thermo Scientific)
- nanoLC System (e.g. Dionex UltiMate 3000 UHPLC, Thermo Scientific)
- 25cm x 75um IonOpticks Aurora Series UHPLC column (IonOpticks; cat. no: AUR2-25075C18A) **Critical: good chromatography is crucial for obtaining high-quality SCoPE2 data. Columns need to have sharp chromatographic resolution, and additionally, need to be able to tolerate the neutralised TMT in genuine SCoPE2 samples. We have had success with these columns. Columns from other suppliers may also work well, but will require testing**.

- In-source blower elbow: Idex Health & Science, part #: P-432
- ABIRD, Active Background Ion Reduction Device (ESI Source Solutions; cat. no: ABFLEX-TM). **Note: recommended if using an ion source that is open to the room, for example the Nanospray Flex Ion Source**.
- nanoLC Column Heater & controller, e.g. Bufferfly heater/controller PST-BPH-20, PST-CHC (Phoenix S&T)

### Software

- MaxQuant Software (v1.6.7 or newer), available at maxquant.org with free registration^46,47^. Other software (e.g., Proteome Discoverer, Comet, and FragPipe) can be used with minor adjustments of DART-ID and DO-MS software to the output of these different search engines. Note, TMTpro 16plex modifications are not included with this release, but an updated modifications.xml file including these forms part of the supplementary material provided with this manuscript.
- DART-ID Software, freely available from dart-id.slavovlab.net/^24^. The DART-ID software allows FDR-controlled and improved peptide identification based on a Bayesian approach that updates peptide PEPs using informative peptide features, e.g. retention times. Other software packages can also enhance peptide identification using informative peptide features^48,49^.
- DO-MS Software, freely available from do-ms.slavovlab.net/^25^. The DO-MS software allows for visualisation of mass spectrometry run features, which is the basis for specifically diagnosis problems and optimizing data acquisition. **Note: For optimal use, the “Peak features” option must be enabled in the MaxQuant analysis options**.

### Reagent Setup

- **Tandem Mass Tags:** Tandem Mass Following Thermo Fisher recommendations, 5 mg of TMT powder should be resuspended in 200 ul of anhydrous acetonitrile, which produces a stock concentration of 85mM. Thermo Fisher recommends storing unused TMT reagents at -20C with desiccant for a period of one week. We have found that storing aliquotted TMT labels at -80° C works as well. **Critical: use of alternate (non-anhydrous) acetonitrile can result in loss of TMT reactivity and poor downstream labelling**.

### Equipment Setup

#### Instrument setup

As no cleanup is performed on SCoPE2 samples, material can rapidly accumulate on the heated capillary during SCoPE2 experiments, requiring cleaning and recalibration on a more regular basis than desirable. Minor modifications to the exterior of the mass spectrometer and run method avoid this by ensuring that material eluting from the LC during sample loading doesn’t enter the instrument.

On Orbitrap instruments, the ion source is used with the ion sweep cone removed. An inexpensive blower elbow is attached to the sheath gas outlet above the heated capillary, and aimed at the capillary outlet. The attachment of this blower elbow to a Q-Exactive-type instrument is shown in Fig. 1. This is used in combination with a modified run method in Xcalibur. This approach switches through several tune files which regulate the spray voltage and sheath gas flow rate. We highlight our LC-MS/MS run method for SCoPE2 samples below illustrating these switches. During the 20 minute loading period, a first tune file supplies no voltage to the source, causing material eluting from the column to collect on the emitter. For the final 20 seconds of loading, a second tune file keeps the voltage off, whilst applying sheath gas. The sheath gas is directed at the emitter tip through the blower elbow attached above, removing the accumulated material. Finally, the method switches to a normal tune method, representing the latest instrument calibration where the voltage is applied and sheath gas turned off. Spectra are only recorded when this final tune file is applied. As no spectra are recorded for the first two tune files, these do not require updating during regular instrument calibration and maintenance.

**Figure 1.**
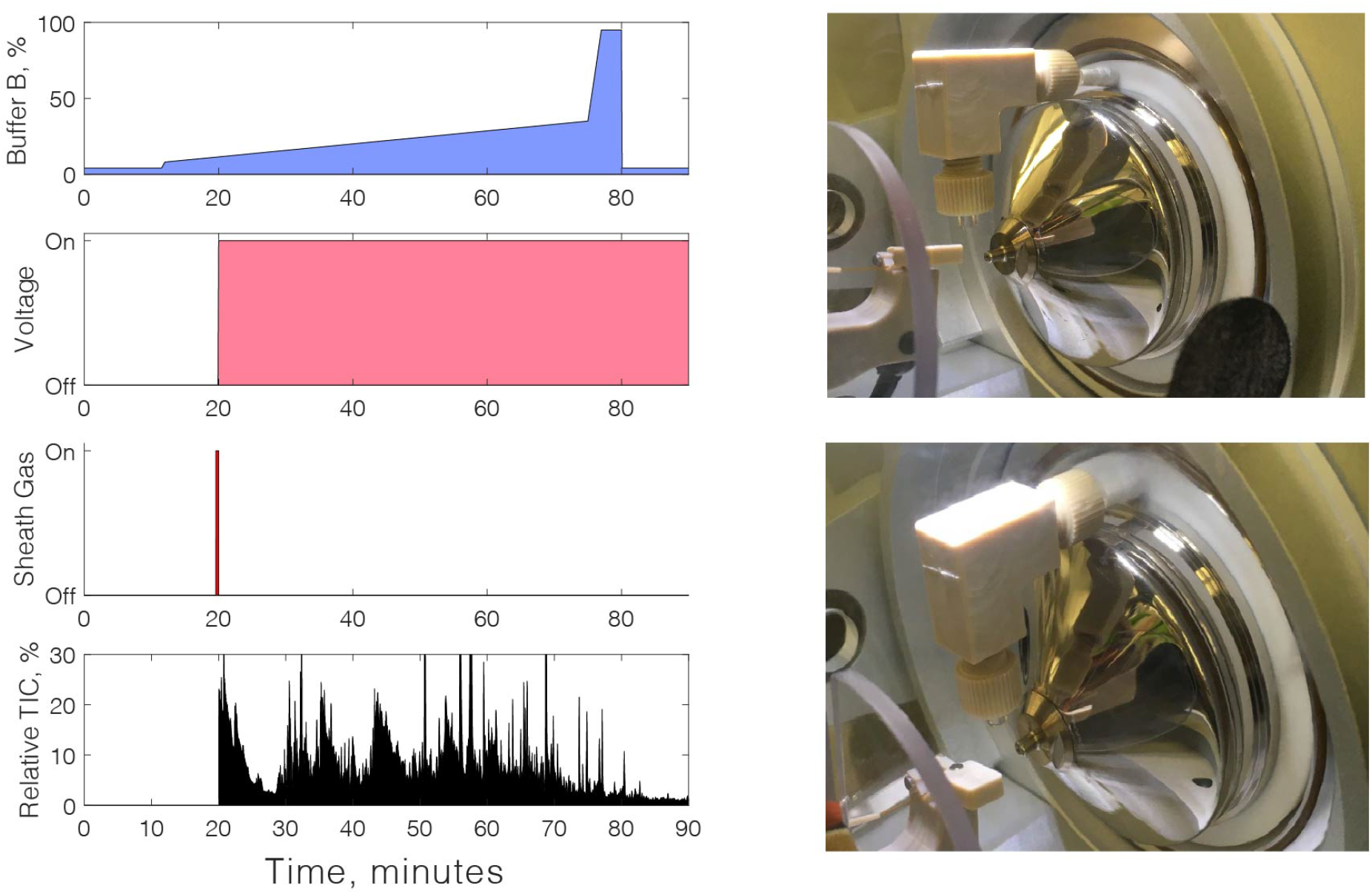
LC-MS/MS setup for SCoPE2 experiments. The left panels show the typical gradient parameters used for SCoPE2 runs. Non-standard portions of the gradient include turning the voltage off during the initial phase to the gradient to reduce the contamination of the heated capillary. Material collected on the emitter tip is then removed with sheath-gas briefly directed at the tip through a blower elbow. After this point the voltage is applied and scan data collected. The right panels show the attachment of the blower elbow to a Q-Exactive classic instrument.

If the laboratory atmosphere has a high background ion levels, these may pose difficulties for low abundance sample analysis, especially with an source open to the room atmosphere such as the flex ion source. In such case, the level of contaminant ions can be significantly reduced by using the ABIRD device, see Extended Data Figure 1. The ABIRD device directs HEPA-filtered air towards the source, thus reducing background ions significantly. The benefits of using an ABIRD are pronounced only when the atmosphere is contaminated.

#### Liquid chromotography setup

nLC performance is crucial for obtaining high quality quantification data from SCoPE2 samples. The key criteria for column selection, is short full-width half-maximum (FWHM) peak widths. However, some commercial columns appear to poorly tolerate the high levels of neutralised TMT and hydroxylamine in the SCoPE2 samples and this should also be assessed. The 25cm x 75 *µm* IonOpticks Aurora series column meets both these criteria, and these columns typically offer a life-time of several months for SCoPE2 sample analysis. The column is enclosed in a nanoLC column heater, with the temperature set to 60°. In our setup, the nLC system is operated without a trapping column since trapping columns can contribute to losses of limited samples such as SCoPE2 samples. The nLC system should be plumbed to minimise dead volumes and transfer lengths since those contribute to peak broadening.

#### Autosampler parameters

The General Settings for the Dionex Ultimate WPS-3000 autosampler are given in Table 1.

**Table 1.** Example settings for performing SCoPE2 on a Dionex WPS 3000 autosampler.

| Parameter | Setting |
| --- | --- |
| Draw Speed | 0.050 |
| Draw Delay | 5.000 |
| Dispense Speed | 2.000 |
| Dispense Delay | 2.000 |
| Dispense To Waste Speed | 4.000 |
| Sample Height | 0.000 |
| Puncture Depth | 8.000 |
| Wash Volume | 50.000 |
| Wash Speed | 4.000 |

#### User-defined Pickup Method

The settings for running a user-defined pickup method on the Dionex UltiMate WPS-3000 autosampler are given in Table 2. This pickup method optimizes sample loading time, and thus overall sample throughput. However, it is not essential and the pre-defined *uLPickup* method can be used. Inject mode is set to *UserProg*. Reagent A is set to autosampler location *G1*. This location in the autosampler must contain buffer A or water.

**Table 2.** Example settings for performing SCoPE2 on a Dionex WPS 3000 autosampler.

| Command | Parameters |
| --- | --- |
| UdpDraw | ReagentAVial, 5[ul], 0.2 [ul/s], 5 [mm] |
| UdpMixWait | 5 [s] |
| UdpDispense | Drain, 0.000, 2 [ul/s], 5 [mm] |
| UdpInjectValve | Load |
| UdpDraw | SampleVial, Sampler.Volume, 0.2 [ul/s], 0 [mm] |
| UdpMixWait | 5 [s] |
| UdpDispense | Drain, 0.000, 2 [ul/s], 5 [mm] |
| UdpDraw | ReagentAVial, 2.4 [ul], 0.2 [ul/s], 5 [mm] |
| UdpMixWait | 5 [s] |
| UdpDispense | Drain, 0.000, 2 [ul/s], 5 [mm] |
| UdpInjectValve | Inject |
| UdpInjectMarker |  |
| UdpMixWait | 5 [s] |
| UdpMixNeedleWash | 200 [ul] |

#### LC-MS/MS run parameters

The run method parameters given in Table 3 are what is used on the authors’ instrument. They represent a good starting point, but may not yield optimal performance on a different system without substantial optimization. Details on how to optimize the run method for the users system are given within the protocol.

**Table 3.** Example settings for performing SCoPE2 on a Thermo Q-Exactive (classic) Instrument. These settings will vary for individual instruments even from the same model, and for different experimental designs. However, these may serve as a convenient starting point for optimization. Below we discuss how to determine specific instrument settings for each use-case.

| Parameter | Setting |
| --- | --- |
| Spray Voltage | 1,800 - 2,500 V |
| Capillary temperature | 250°C |
| Full-scan MS range | 450-1600 m/z |
| MS1 resolution (m/z 200) | 70,000 |
| Maximum injection time | 100 ms |
| AGC target | 1e6 |
| TopN | 7 |
| Precursor charge state | 2-5+ |
| (N)CE | 33 |
| MS2 resolution (m/z 200) | 70,000 |
| MS2 accumulation time | 300 ms |
| MS2 AGC target | 5e4 |
| MS2 AGC minimum | 2e4 |
| Isolation width | 0.7 m/z |
| Isolation offset | 0.3 m/z |
| Charge exclusion | Unassigned, 1, $\geq 4$ |
| Dynamic exclusion | 30 s |

These parameters in Table 3 may need a slight adjustment for the new line of Orbitrap instruments, such as Exploris and Eclipse. A primary advantage of the newer Orbitraps is that they need less transient time to offer the high resolution needed by SCoPE2 (60,000 – 70,000). This advantage does not change significantly the data acquisition parameters, because the speed of SCoPE2 analysis is usually limited by the time needed for ion accumulation rather than the transient time needed to achieve high resolution^18,19^. Thus, we expect that most parameters outlined in Table 3 remain a good starting point for the newer Orbitrap instruments. With Exploris and Eclipse, we recommend doubling the resolution for MS2 scans because such an increase of the resolution increases the signal-to-noise ratio at no cost^18^, that is it does not slow down the rate of acquiring MS2 scans. These theoretical expectations have been consistent with results from our collaborations with colleagues using the latest Exploris and Eclipse Orbitrap instruments. We have recently begun performing SCoPE2 analysis with a timsTOF system, and our preliminary data show much promise of the system. The higher sensitivity of the TOF detector reduces the size of the isobaric carrier needed to achieve the designed tradeoff of throughput and copy number sampling from the single cells^19^. We hope that our parameters and guidelines are a useful starting point, and we recommend that each laboratory optimize their mass spectrometry systems for single cell analysis, as each instrument might need slightly different parameters.

Gradient details are found in Table 4. Typical gradients and the voltage and sheath-gas switching can also be found in Figure 1.

**Table 4.**
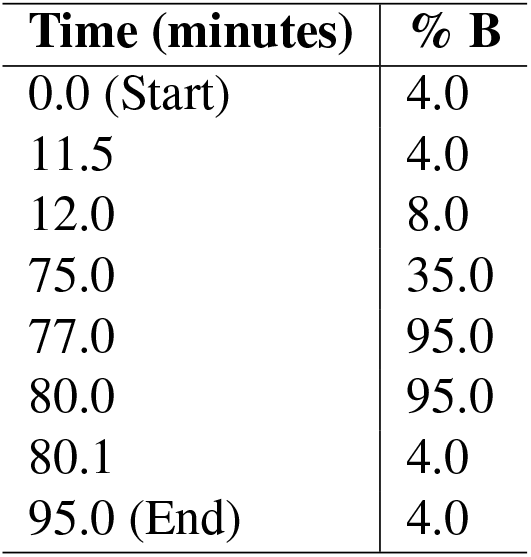
Example LC gradient for performing SCoPE2 on a Thermo Q-Exactive (classic) Instrument with a Dionex 3000 nLC. Further gradient optimization may be required depending on the use case. Buffer A is 0.1% formic acid in HPLC-grade water, Buffer B is 0.1% formic acid in 80% acetonitrile/20% HPLC-grade water. The gradient is run without a trap column at a constant 200 nL/minute flow rate.

| Time (minutes) | % B |
| --- | --- |
| 0.0 (Start) | 4.0 |
| 11.5 | 4.0 |
| 12.0 | 8.0 |
| 75.0 | 35.0 |
| 77.0 | 95.0 |
| 80.0 | 95.0 |
| 80.1 | 4.0 |
| 95.0 (End) | 4.0 |

#### Mantis liquid dispenser setup

The liquid handler will require calibrating for the 384-well plates used for preparing SCoPE2 samples, as per the instruments normal method. 384-well and 96-well plates benefit significantly from use of the PCR plate adaptor (see equipment), and we strongly recommend its use. A practice dispense of 1*µ*l of water, in a spare plate prior to using the system for SCoPE2 processing is recommended to visually confirm plate alignment and liquid handler calibration prior to use.

### Software setup

#### MaxQuant

MaxQuant can be freely downloaded from *maxquant*.*org*, following free registration. For users unfamiliar with standard MaxQuant usage, an annual summer school is offered. Videos from past years can be found on YouTube, and documentation is linked to from *maxquant*.*org*.

Where search parameters used for SCoPE2 analysis deviate from the MaxQuant defaults, these are given in Table 5. There are two profiles of settings used for analysis of SCoPE2 data with MaxQuant. The first (listed under Variable search parameters), is used to determine labelling efficiency with TMT or TMTpro reagents. The second is for searching SCoPE2 data or 100xMaster data. Custom modifications are required for the variable search and are included with the *modifications*.*xml* file that forms part of the supplementary material. This should be used to replace the *modifications*.*xml* file that comes with MaxQuant and can be found in the MaxQuant directory, in */bin/conf/*. If you wish to generate these custom search modifications yourself, simply duplicate the TMT or TMTpro 126 N-terminal and Lysine modifications and change the label type from isobaric label to standard. If using TMTpro reagents, MaxQuant 1.6.7 does not come with these in its library and they have been included in the *modifications*.*xml* file. A *mqpar*.*xml* file is also included that can be loaded into MaxQuant which pre-loads TMTpro isobaric labels.

**Table 5.** Maxquant settings. MaxQuant adjustment from default settings for using MaxQuant to assess TMT labelling efficiency (Variable search) or for using MaxQuant to analyse SCoPE2 or 100xMaster data. † TMTpro modifications are not included in the 1.6.7 release of MaxQuant. If wishing to search TMTpro 16plex data, a modifications.xml file and mqpar file to load the TMTpro reagents into Maxquant 1.6.7 is included in supplementary data. *custom modifications named as per the included modifications.xml file.

| Parameter name | Setting |
| --- | --- |
| **Variable search parameters** |  |
| *Group-specific parameters* |  |
| → *Type* | Standard |
| → *Variable Modifications (All)* | Acetyl (K) |
|  | Acetyl (N-term) |
| → *Variable Modifications: 11plex ONLY* | Variable_TMT10plex N-term* |
|  | Variable_TMT10plex Lys* |
| → *Variable Modifications: 16plex ONLY†* | Variable_TMTpro16plex N-term*† |
|  | Variable_TMTpro16plex Lys*† |
| *All other parameters default* |  |
| **SCoPE2 search parameters** |  |
| *Group-specific parameters* |  |
| → *Type* | *Reporter ion MS2: 11plex TMT or 16plex TMTpro†* |
| → *Variable Modifications* | Oxidation (M) |
|  | Acetyl (Protein N-term) |
| → *Fixed Modifications* | None |
| *Global parameters* |  |
| → *Identification* | PSM FDR: 1.00 (if using DART-ID) |
|  | Protein FDR: 1.00 (if using DART-ID) |
| → *Advanced identification* | Uncheck *Second peptides* |
| → *Advanced* | Check *Calculate Peak Properties* |
| *All other parameters default* |  |

The isobaric tags of TMT contain isotopic impurities, which affect SCoPE2 quantification as they do all TMT workflows. Because of the impurities, a portion of each sample is labeled with a tag whose mass is different (mostly by ±1 Da) of the intended mass of the tag. The influence of such isotopic impurities can be modeled with a linear superposition model and computationally mitigated by performing a deconvolution. This is a standard part of many workflows, including the MaxQuant one, and we recommend inputting the TMT batch-specific impurities supplied by the manufacturer for achieving optimal correction. With MaxQuant, this option can be accessed via the “Group-specific parameters” and then the “Type” tab, Reporter ion MS2 will first need to be selected. Then, the isotope correction factors can be imported directly into the software, using the “Import” button. In the context of isobaric carriers, the impurities have an additional impact: The impurities of the tag used for the abundant carrier peptides may have a disproportionate impact on a couple of neighbouring channels, and thus we recommend not using these channels for single-cell samples.

#### DART-ID

The DART-ID^24^ project website which contains detailed installation and usage instructions can be found at dart-id.slavovlab.net/.

#### DO-MS

The DO-MS^25^ project website which contains detailed installation and usage instructions can be found at do-ms.slavovlab.net/.

### Procedure

We recommend the use of the newer TMTpro 16plex reagents for their higher throughput and reduced cost per single cell. If you wish to use 11plex reagents sections where the protocol deviates will be indicated. **11plex**.

### Generation of 100xMaster and 100xCarrier Samples Timing: 1-2 days

**Critical: refer to Table 6 for recommended labelling strategy for 100xMaster and 100xCarrier samples**.

**Table 6.** Recommended 100xMaster and 100xCarrier-only sample design. Master standards diluted to 1x allow convenient testing and optimisation of the LC-MS/MS system without the variability seen in true single cell samples. The use of a carrier-only sample allows testing for sample carryover in the single-cell channels, and detection of low-level contaminants which could not be identified by using ‘blank’ runs. The table lists **11plex** Tandem Mass Tag reagents, and the section from 128C to 131C could be extended if using **16plex TMTpro** reagents, keeping the layout for 126-131C and adding additional 100 cells, cell type A or B in 132N-134N.

| 100xMaster | 100xCarrier | TMT label |
| --- | --- | --- |
| 5000 cells, cell type A | 5000 cells, cell type A | 126 |
| 5000 cells, cell type B | 5000 cells, cell type B | 127N |
| Unused | Unused | 127C |
| Unused | Unused | 128N |
| 100 cells, cell type A | Unused | 128C |
| 100 cells, cell type B | Unused | 129N |
| 100 cells, cell type A | Unused | 129C |
| 100 cells, cell type B | Unused | 130N |
| 100 cells, cell type A | Unused | 130C |
| 100 cells, cell type B | Unused | 131N |
| Unused | Unused | 131C |

**Table 7.** utomated and manual options for performing the SCoPE2 protocol. Placing the 384-well plates with sorted cells into the liquid handlers and the pooled SCoPE2 samples into the autosamplers is performed manually. The last column indicates the corresponding procedure steps numbers in the main text.

| Protocol Step | Automated options | Manual options | Step No. |
| --- | --- | --- | --- |
| Cell Isolation | FACS or CellenONE | Manual cell picking<sup>†</sup> | (22 - 25) |
| Cell lysis | Liquid handler (e.g., Mantis) | Multi- or single channel pipette | (40 - 42) |
| Protein digestion | Liquid handler (e.g., Mantis) | Multi- or single channel pipette | (43 - 46) |
| TMT Labeling | Liquid handler (e.g., Mantis) | Multi- or single channel pipette | (48 - 54) |
| Pooling samples | Not used | Single Channel pipette | (55 - 57) |
| Loading samples | Autosampler | Not used | (65) |
| LC-MS/MS | Xcalibur or timsControl | Not used | (65) |

1. Obtain single-cell suspensions of two mammalian cell lines (thereafter termed samples A, B) of interest in ice-cold PBS, from the same species (e.g. for human cells, U-937 and Jurkat). **Note: the exact means by which these cell lines are prepared as single-cell suspensions will vary depending on the cell lines of interest**.
2. Count the cells (a hemocytometer is sufficient), and resuspend 40,000 of each cell type in separate PCR tubes in a volume of 20*µ*l of HPLC-grade water, resulting in 2,000 cells/*µ*l. **Note: Controlled experiments have demonstrated that the freeze-heat lysis method, named mPOP, is as efficient as classical urea lysis**^**20**^. **We have also performed BCA protein content assays on different cell types lysed via mPOP. The BCA data confirmed the estimated protein levels per single cell between two technical replicates per cell type. HeLa cells are among the largest cell types: We extracted 0.5 ng of protein per cell, which agree with the expected total protein per HeLa cell**^**50**^. **Additionally, Jurkats were among the smallest cells we tested, which was confirmed by the scaled protein content, of around 0.1 ng of protein per cell. Therefore, if we extract protein from 40**,**000 HeLa cells, we would expect a protein yield of around 20**,**000 ng; likewise, from 40**,**000 Jurkats, we would expect 4**,**000 ng**.
3. Freeze PCR tubes containing cells at -80°C for a minimum of 30 minutes. **Note: this represents a potential pause point in the protocol**.
4. Transfer PCR tubes to a thermocycler with a heated lid, and heat to 90°C for 10 minutes (set lid to 105°C), holding the sample at 12°C when the heating cycle has completed.
5. Once the samples have cooled to 12°C, add 1*µ*l of benzonase nuclease diluted to ≥ 5 units/*µ*l.
6. Briefly vortex the samples for 3 seconds, followed by centrifugation in a bench-top PCR tube spinner for 3 seconds to collect liquid.
7. Sonicate the samples for 10 minutes in a water bath sonicator, and transfer to ice.
8. Once on ice, supplement both samples with 4*µ*l of a mastermix containing:
9. Briefly vortex the tubes to mix, followed by brief centrifugation to collect liquid.
10. Transfer the samples to a thermocycler and digest for 3 hours at 37°C (Heated lid set to 52°C).

11. **For 100xMasters:** add 3.2*µ*l of sample A to each of 4 PCR tubes (1 for the carrier channel, 3 for the ‘single-cell’ channels). Do the same with 4 new PCR tubes for sample B. **For 100xCarrier:** add 3.2*µ*l of sample A to 1 PCR tube. Do the same with a new PCR tube for sample B. **Note: this step can be done with larger volumes of cells. Adjust the volume of subsequent reagent additions proportionally to the volume of the input digested sample**.
12. Following the layout detailed in Table 6, add 1.6*µ*l of 85mM TMT label to each of the tubes. E.g. for a 100xMaster, the four PCR tubes containing sample A should be labelled with 126, 128C, 129C and 130C.
13. Briefly vortex the tubes, spin down to collect liquid.
14. Incubate tubes at room temperature for 1 hour.
15. Add 0.7*µ*l of 1% HA to each tube.
16. Briefly vortex to mix, followed by centrifugation to collect liquid.
17. Incubate tubes at room temperature for 30 minutes.
18. **For 100xMasters:** combine both carrier sample tubes (126, 127N) in a single glass insert. Dilute the remaining samples comprising the ‘single-cell’ samples to 25*µ*l and add 0.5 of each to the glass insert. This produces dilutions for the carrier and ‘single-cell’ samples as described in Table 6. Mix well by pipetting **For 100xControls:** Combine both tubes into a glass insert, mixing well by pipetting.
19. Divide the 100xMaster or 100xControl samples into 5 aliquots of equal volume. These each contain 20 injections of material. **Note: if different aliquot sizes are required, adjust the subsequent resuspension volumes proportionally, keeping the volume at 1***µ***l per injection**.
20. Use a speedvac to reduce the samples to dryness. These can be stored dry at -80°C until required.
21. To use an aliquot, resuspend in 20*µ*l of 0.1% formic acid. See the following section for guidance of LC-MS/MS optimization (? Troubleshooting)

| Component | Amount | Final Concentration per tube |
| --- | --- | --- |
| Trypsin Gold (200 *ng*/μl) | 1.3μl | 10ng/μl |
| Triethylammonium bicarbonate (TEAB) pH 8.5 | 2.5μl | 100 *mM* |
| HPLC-grade water | 0.2μl | Not applicable |

### LC-MS/MS optimization using 100xMaster Samples Timing: 1+ days

**Note:** While it is possible that optimization could be accomplished in a couple of days with a well-running LC-MS/MS system set up as described in instrument setup, deviation from this setup may result in significant increases in the amount of time required for instrument optimisation.

The key elements to optimize are efficient sample delivery to the mass spectrometer and sampling peptide-like ions as close to their elution peak apex as possible. The latter both maximizes signal intensity and minimizes co-isolation with other ions which can compromise MS2-based quantitation using TMT. While a number of software packages can be employed to help optimize these parameters, we recommend using MaxQuant and DO-MS as described previously^25^. In order to generate all plots within the DO-MS dashboard, users must set up MaxQuant using the parameters found in Table 5.

#### Optimizing data acquisition

##### Motivation

Given that input amounts in SCoPE2 samples are lower than for typical bulk proteomics preparations, optimizing chromatography to maximize the number of ions of each species delivered per unit time is of primary importance.

- **Sample Delivery Optimization**
  ‐ By plotting peak widths, both at base and at full-width half max, this consideration can be evaluated. Optimizing peak widths involves considerations of gradient length, gradient steepness, column chemistry, and LC plumbing, among others.
- **Optimizing Sampling at the Elution Peak Apex**
  ‐ There are several interrelated instrument parameters that directly impact ion sampling: MS2 fill times, top N, and AGCmin. By adjusting the length of time that the instrument collects ions at the MS2 level and changing the number of ion species that are selected for MS2 analysis, it is possible to alter the point in the elution profile where the ion is sampled.
  ‐ Establish an Apex Offset performance baseline by running two replicate 1x injections of the master sample
  ‐ Search these 1x injections using MaxQuant (sample MaxQuant parameters provided in the Supplemental Data section)
  Load the MaxQuant output into DO-MS, and navigate to the Ion Sampling dashboard tab.
  ‐ The Apex Offset plot window presents the temporal distance between when each ion was selected for MS2 analysis and that ion’s elution apex, when it was most abundant. In general, it is preferable to bias MS2 sampling of an ion as close to its elution apex as possible.
  ‐ If the Apex Offset distribution is biased towards early sampling, increase the length of the duty cycle by increasing MS2 fill times or increasing the number of ions selected for MS2 analysis (top N). If the Apex Offset distribution is biased towards sampling ions after their elution apices, decrease the MS2 fill time or top N. In practice, it will not be possible to target all peptide-like ions in a complex mixture at their elution apices.
- **Assessing Coisolation:** A straightforward metric for assessing MS2 spectral purity, or the degree to which coisolation of multiple precursors prior to precursor fragmentation is occurring, is generated by MaxQuant as Parent Ion Fraction (PIF). A distribution of PIF values on a per-experiment basis can be found in DO-MS’ Peptide Identification tab.

#### Carrier Channel Assessment

##### Motivation

- It is important to determine the degree of labeling of the carrier channel peptides, as any unlabeled peptides in this channel could be labeled by unquenched single-cell sample barcodes upon pooling of the carrier channel, reference channel, control well samples, and single-cell samples.
- Before combining the bulk prepared carrier channel with your TMT-labeled single-cell samples, a 1x injection of the carrier material should be run separately to assess labeling efficiency, as well as missed cleavage rate.
- If only 1% of the peptides in the carrier channel are subject to this cross-labeling, then the unlabeled material is equivalent to the peptide input of a single-cell. Labeling efficiency of the carrier channel can be assessed using a custom-defined modification within MaxQuant, shown in Table 5 under Variable search parameters. These modification accounts for the mass of a TMT label on either the N-terminus or any lysine residues present in a given sequence, and will allow the fraction of labeling sites that were successfully labeled to be determined. The results are quantified and visualized by a dedicated tab of DO-MS.
- Additionally, a high number of missed cleavage sites in the carrier channel input indicates suboptimal digestion conditions and can impair peptide identification.
- If either the labeling efficiency is low (at least 99 percent labeled) or the missed cleavage rate is high (less than 15 percent), a new carrier channel should be prepared if possible. Relabeling the carrier channel is often not a successful course of action due to the presence of hydroxylamine from the previous quenching reaction.

#### Metrics for evaluating single-cell quantification

##### Motivation

Many factors may undermine data quality, including the possibility of inefficient cell isolation (sorting), poor protein digestion, incomplete TMT label quenching, low TMT reactivity due to partial hydrolysis, and background contamination. Thus, we strongly recommend using built-in controls to benchmark SCoPE2 data quality.

- The distributions of reporter ion intensities (RII) for each label, normalized by those of the most abundant sample’s label, will allow relative estimates of the peptide input in each channel. For instance, the median intensity value in the distributions of relative reporter ion intensities (RRII) in the single-cell channels should be one-hundred-fold less than the normalized median RII in the carrier channel, if the carrier channel contains digested peptides from 100 cells. Channels which contain median RRII equivalent to those found in the control wells may be failed wells. Channels which contain median RRII equivalent to the median RRII in the reference channel may not have been properly quenched, allowing for cross-labeling of abundant material from the carrier channel during sample combination. A plot of RRII distributions can be found in DO-MS’ Single-Cell Diagnostics tab.
- A correlation matrix of reporter ion intensities is another useful metric that often can detect major problems in sample preparation. After removing peptide spectral matches below a chosen confidence cut off, reverse matches, and contaminants from the experimental data set, the pairwise correlations between vectors of RIIs for each channel, displayed as a correlation matrix, is useful for assessing whether cross-labeling has occurred. In general, the RIIs for cells of one type should correlate more highly with one another than with cells of another type or control wells. Such a correlation matrix can be found in DO-MS’ Single-Cell Diagnostic tab.

### SCoPE2 sample preparation Timing: 1-2+ days

Preparation of SCoPE2 sets can be divided into three key stages: i) Initial preparation of 384-well plates containing sorted single cells and control wells for the system under investigation. ii) Preparation of the carrier and reference channel material in bulk, and iii) Preparation of the 384-well plates containing single-cell material, and its combination with the carrier and reference material to generate completed SCoPE2 sets. A single 384-well plate, along with its carrier and reference material can be sorted and processed in 1-2 days. Additional 384-well plate can be processed faster with about two hours of hands-on time per plate.

### SCoPE2 sample preparation part I: Isolating single cells Timing: 1h 20m +

Isolating single cells can be accomplished by fluorescence-activated cell sorting (FACS) or by CellenONE as demonstrated in this protocol. When using CellenONE, samples can be prepared by nPOP as described in ref.^21^. We have successfully used Aria II, Aria III and Sony MA900 sorters. Other single-cell dispensers, such as Namocell, might also be usable, but we do not have direct experience with them.

Ideally, single-cell isolation / sorting should be evaluated by simple and direct means, e.g., by using colorimetric assays. Its success should be maximized by ensuring stable spray and good alignment for FACS sorters and low static electricity for CellenONE. If the isolation process is not well validated, the use of positive control well (containing cell lysate diluted to single-cell level) is strongly encouraged.

22. Add 1 *µ*l of HPLC-grade water, supplemented with 25 *fmol* per peptide per *µ*l of Waters MassPrep to each well of a 384-well PCR plate. Seal the plates and then centrifuge for 20 seconds (subsequent use of a plate spinner in this manner will be marked **spin down**, and can either be used immediately or stored at -20°C until required). **Note: while the use of a liquid dispensing robot is strongly recommended, it is not essential for this step. Note: this represents a pause point in the protocol**.
23. Prepare a disaggregated suspension of unfixed cells, washed twice with 1x ice-cold phosphate buffered saline (PBS), and resuspended in ice-cold PBS should be prepared. Exactly how this is achieved will vary depending on the cell type of interest. Some examples are given below: **Suspension cells:** Centrifuge to remove cell culture media, and wash with, and then resuspend in ice-cold 1x PBS. **Adherent cells:** Trypsinise or remove cells from a dish surface by scraping. Centrifuge to remove cell culture media. Then wash with, and resuspend in ice-cold PBS.
24. Distribute single cells into the 384-well plates prepared earlier. A suitable method is the use of a cell sorter, with care to ensure that the sorted population represents single live cells of interest. **Critical: As described in the section on experimental design, the distribution of particular cells on and between different 384-well plates should include no-cell controls (negative control samples). If multiple cell treatments are analyzed, these should be distributed equally across the plates to avoid pairing cell types and batch effects**. **Note: the wells on the outer edge of the 384-well plate may experience edge effects during heating and incubation steps leading to more rapid evaporation. You may wish to omit these wells from use**.
25. After isolating single cells into the plates, seal and **spin down** the plates, and then transfer to a -80°C freezer as rapidly as possible. **Note: this represents a pause point in the protocol. Ideally all the cells of interest should be prepared in a single batch**.

### SCoPE2 sample preparation part II: carrier and reference channel processing Timing: 1 day

The number of cells used for isobaric carrier can be determined based on the principles and trade offs established from controlled experiments^19^. For the default size of 100-200 cell carrier, you need about 11,275 cells to prepare the carrier and the reference per 384-well plate and fewer cells can be used of needed. If cells are not limited, we recommend retaining about 22,000 cells per 384-well plate being prepared. This allows for a 200 cell carrier, 5 cell reference channel per SCoPE2 set, and also minimises dilution or small volume pipetting steps. Cells should be combined according to the carrier/reference design, washed in PBS, then resuspended in 11*µ*l of HPLC-grade water in a PCR tube and stored at -80°C until the **carrier and reference channel processing** stage of the protocol. **Critical: it is recommended that carrier material, and absolutely essential that reference material, be prepared as a single batch, sufficient for the number of 384-well plates being prepared for the SCoPE2 experiment. The numbers given here detail preparing sufficient carrier/reference material to prepare a single 384-well plate of SCoPE2 samples and should be scaled up accordingly for increased numbers of plates**.

26. Remove the PCR tube containing the sample from the -80°C freezer, and transfer to a PCR machine as rapidly as possible.
27. Heat the sample to 90°C for 10 minutes (heated lid at 105°C), then allow it to cool to 12°C, and **spin down** using a PCR tube spinner.
28. Transfer the sample to a water bath sonicator, and sonicate for 5 minutes at room temperature, and **spin down**.
29. To the sample, add 2.2*µ*l of HPLC-grade water containing the following: 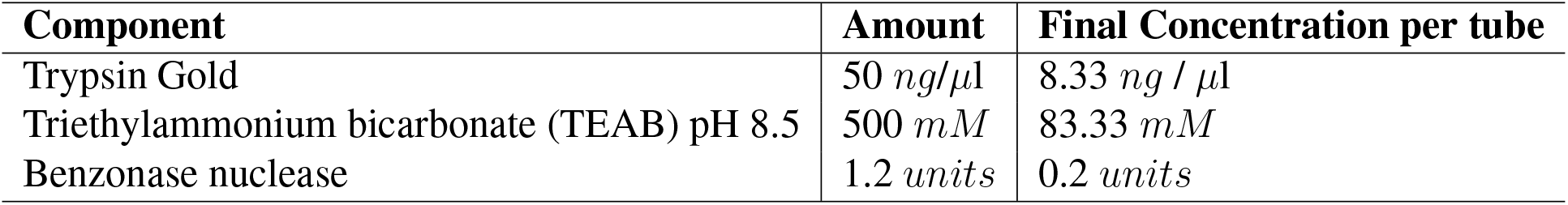
30. The sample PCR tube should be vortexed, and **spun down**.
31. Incubate the sample in a 96-well PCR machine at 37°C (heated lid 52°C) for 3h.
32. After digestion, **spin down** the sample, and split into two equal 6.6*µ*l volumes each containing 11,000 cells in separate PCR tubes.
33. To one sample, add 3.3*µ*l of 126 TMT label (85 *mM*). This will become the **carrier** material. To the other sample add 3.3*µ*l of 127N TMT label (85 *mM*). This will become the **reference** material. This step is identical whether **11plex TMT** or **TMTpro 16plex** reagents are used. In both cases, the 126 and 127N labels are used.
34. **Spin down** the two samples and incubate at room temperature (22°C) for 1 hour.
35. Add 1.65*µ*l of 0.5% hydroxylamine diluted in HPLC-grade water to each of the two samples. Then vortex the tubes and **spin down**.
36. Incubate the tubes at room temperature (22°C) for 30 minutes.
37. Dilute the 127N-labelled tube containing the **reference** material by taking 2*µ*l of the reference and mixing it with 78 *µ*l of HPLC-grade water. Then the carrier and diluted reference material can be mixed 1:1 and subsequently further diluted so that carrier and reference concentrations are 100-200 cells/*µ*l and 5 cells/*µ*l, respectively. (For example: combine 8*µ*l of carrier with 8*µ*l of diluted reference, then add 22*µ*l of HPLC-grade water to obtain 200 cells/*µ*l carrier and 5 cells/*µ*l reference.) **Note: If preparing more than one 384-well plate as part of a SCoPE2 experiment, it is recommended to prepare carrier and reference material in bulk, aliquot the carrier and reference channel material in SCoPE2 set aliquots, so they can be used per 384-well plate. The volumes described in this section can be scaled up relative to cell numbers**.
38. Vortex and **spin down** the combined **carrier/reference** material. Store at -80°C until required in the following section. Consider not diluting the mixed carrier and reference material if not using right away. **Note: this represents a pause point in the protocol**.
39. Assess the quality of the carrier material prior to combining it with single cell samples in the following section III. Such quality checks include miscleavage rate and labeling efficiency, which should be less than 15 percent and greater than 99 percent, respectively. Additionally, overlabeling should be assessed. This can be done in MaxQuant, by specifying TMT Pro labels as a variable modification on histidine, serine, threonine, and tyrosine, and as a fixed modifications on primary amines (N-terminus and lysine). (? Troubleshooting) **Note: We have found that overlabeling is not a major issue in this protocol. In the data provided with this paper, overlabeled peptides account for less than 3 percent of all confidently identified peptides**.

### SCoPE2 sample preparation part III: single-cell processing SCoPE2 set generation Timing: 8h (2h hands-on) per 384-well plate

**Note: For the following steps, a single 384-well plate should be prepared at a time**.

40. Remove the 384-well plate from the -80°C freezer and transfer to a 384-well PCR machine as rapidly as possible.
41. Set the PCR machine to heat the 384-well plate to 90°C for 10 minutes with the heated lid set to 105°C, followed by cooling the plate to 12°C. **Spin down** the plate.
42. Transfer the 384-well plate to a water bath sonicater. Sonicate for 5 minutes at room temperature, and **spin down**.
43. While sonication is being performed, prepare 100 *µ*l of a mastermix per 384-well plate, which contains the following:

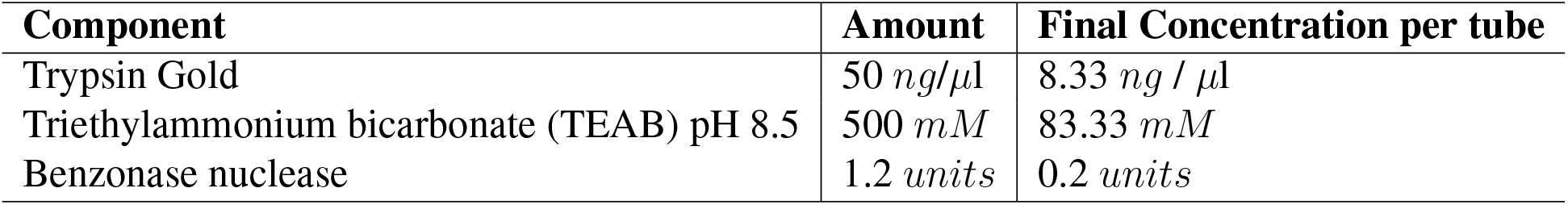 **Note: the use of alternative sources of trypsin requires optimisation, as some suppliers have higher levels of contamination and perform poorly for SCoPE2**.
44. Using a liquid handler (e.g. Mantis), dispense 0.2*µ*l of mastermix into each well of the 384-well plate. **Note: if using manual liquid dispensing, use of larger volumes is recommended to reduce handling errors**.
45. After adding the mastermix to all wells of the 384-well plate, seal the plate, vortex for 5 seconds and **spin down**
46. Incubate the 384-well plate in a 384-well PCR machine at 37°C, with the heated lid set to 52°C for 3 hours.
47. After digestion, **spin down** the plate.
48. As illustrated in Table 8, a typical layout for a SCoPE2 experiment has a 126-labelled carrier channel, 127N-labelled reference and 127C left blank. This leaves 8 channels available for labelling single cells or control wells from 128N-131C. If using TMTpro 16plex reagents, the SCoPE2 layout remains similar, and is shown in Table 9. **Critical: The placement of carrier, reference and empty channels in these designs helps prevent contamination of single-cell data due to isotopic contamination of these TMT labels from the much more abundant carrier and reference channels. The extra empty channel included with the 16plex TMTpro is due to the current higher level of isotopic contamination with these reagents compared with 11plex TMT**.

**Table 8.** SCoPE2 Sample Labelling Strategy. **11plex TMT** labels are highlighted in **bold**. 126 and 127N are used for the carrier and reference which are prepared in bulk in the previous **carrier and reference channel processing** portion of the protocol and diluted. 128N-131C are used for labelling SCoPE2 single-cell and control samples. 127C is unused due to isotopic contamination from 126.

| **126** | **127N** | **127C** | **128N** | **128C** | **129N** | **129C** | **130N** | **130C** | **131N** | **131C** |
| --- | --- | --- | --- | --- | --- | --- | --- | --- | --- | --- |
| Carrier | Reference | Empty | SC | SC | SC | SC | SC | SC | SC | SC |

**Table 9.** SCoPE2 Sample Labelling Strategy using **16plex TMTpro** reagents. 16plex TMTpro labels are shown in **bold**. Sample layout and isobaric reagent usage is essentially the same as when using 11plex TMT reagents. **131N** to **133C** are not shown, but can also be used for single-cell or control samples. The higher level of isotopic carryover from 127N with some TMT reagent lots means that 127C and 128N are both unused due to isotopic contamination from 126 and 127N. However, the total number of single-cell channels that can be used per SCoPE2 set increases from 8 to 12 in comparison with 11plex reagents.

| **126** | **127N** | **127C** | **128N** | **128C** | **129N** | **129C** | **130N** | **130C** | ... | **134N** |
| --- | --- | --- | --- | --- | --- | --- | --- | --- | --- | --- |
| Carrier | Reference | Empty | Empty | SC | SC | SC | SC | SC | ... | SC |

**Table 11.** Protocol Timings. *Steps 5 and 6 will be repeated multiple times as required to achieve the desired experiment depth. Up to 48 SCoPE-sets can be prepared per 384-well plate using 11plex TMT reagents, or 32 SCoPE2-sets with 16plex TMTpro depending on the exact experimental design. The 85 *mM* stocks of TMT labels from 128N-131C should be removed from the -80°C freezer, warmed to room temperature and diluted in anhydrous acetonitrile to 22 *mM*.
49. Using a liquid dispensing robot (e.g. Mantis), add 0.5*µ*l of the diluted TMT reagents to the single cells using the 3PFE chips suitable for high solvent concentrations. **Note: Ensure equal distribution of the different TMT tags so that sets contain one each of samples labeled from 128N to 131C. These can for example be prepared in columns on the 384-well plate**.
50. After adding the TMT reagents to all wells of the 384-well plate, seal the plate, vortex for 5 seconds, and **spin down**.
51. Incubate the plate at room temperature (22°C) for 1 hour.
52. Add 0.2*µ*l of 0.5% hydroxylamine diluted in HPLC-grade water to each well of the 384-well plate using the liquid dispensing robot.
53. After adding the 0.5% hydroxylamine to all wells of the 384-well plate, seal the plate, vortex for 5 seconds, and **spin down**.
54. Incubate the plate at room temperature (22°C) for 30 minutes.
55. Remove the combined carrier and reference material prepared in the previous **carrier and reference channel processing** step from the -80° freezer.
56. For each SCoPE2 set, take 1*µ*l of the carrier/reference material and add it to the first single-cell well of the 384-well plate (128N).
57. Pass the combined carrier/reference material through each of the remaining single-cell wells (128C-131C) to generate a complete SCoPE2 set containing carrier, reference, and 8 single-cell or control samples. **Critical: Combining the samples in this way helps to minimise losses when handling the single-cell material, as the much more abundant carrier material will disproportionately experience the losses**.
58. Take 5*µ*l of 50% acetonitrile prepared in HPLC-grade water and pass it through the same single-cell wells, combining with the carrier/reference/single-cell material in the final well. **Note: This step can help recover any remaining material from the single-cell wells**.
59. Transfer SCoPE2 sets to individual glass autosampler inserts.
60. Place the autosampler inserts into a speedvac, and dry down completely.
61. Once dried, if not running immediately, store the dried SCoPE2 sets at -80°C until required. **Note: this represents a pause point in the protocol**.
62. When ready for LC-MS/MS analysis, resuspend SCoPE2 sets in 1.2*µ*l of 0.1% formic acid in HPLC-grade water.
63. Place the glass autosampler inserts in glass autosampler vials, and capped using screw-thread caps.
64. Vortex the capped autosampler vials for 5 seconds, then centrifuge briefly in a vial spinner to collect the sample at the bottom of the glass autosampler insert. Visually inspected to ensure the sample is at the bottom of the insert, and then place into the autosampler.
65. Inject 1*µ*l of each SCoPE2 set for LC-MS/MS analysis. (? Troubleshooting) **Critical: Instrument and LC method parameters for analysis of individual SCoPE2 sets should be as determined in the previous section on LC-MS/MS optimization using 100xMaster Standards**. **Note: only 1***µ***l of the 1.2***µ***l sample is injected to allow for potential evaporation, or inefficient sample pickup by the autosampler**.

| Section | Time |
| --- | --- |
| **Preparatory Work** |  |
| 1. Production of 100x Master & 100xCarrier Samples | 1-2 days |
| 2. System suitability & Optimization | 1+ days |
| **Single Cell Work** |  |
| 3. Plate preparation | 20 min |
| 4. Cell isolation | 1+ h |
| 5. Sample preparation* | 8 h (2 h hands-on) per 384-well plate |
| 6. LC-MS/MS analysis* | 95 minutes per TMT set |
| 7. Initial data analysis | 1+ days |

### Raw data processing Timing: 1+ hours

66. Search MS spectra for quantified peptides. Raw files corresponding to the SCoPE2 sets constituting a SCoPE2 experiment can be analyzed using a variety of proteomics search engines. For the purpose of accessibility and simplicity, we will focus on settings appropriate to searching the data with the freely-available software MaxQuant. All parameters are specified in the search parameters table.
66. **Optional:** Use additional features to enhance MS data interpretation. The SCoPE2 pipeline uses the DART-ID software to update the confidence of peptide identification, e.g., update the PEPs in the *evidence*.*txt* file generated by MaxQuant. The DART-ID software uses a rigorous Bayesian model to incorporates retention time information in evaluating confidence of peptide identification^24^. For a more complete explanation of the DART-ID software and its usage, please see: https://dart-id.slavovlab.net. Once installed, DART-ID can be run by editing the default configuration file provided to include the path to your evidence file and the path to your desired output location, example:

~~~
input:
 - /path/to/your_search_results/evidence.txt
output:
- /path/to/output_folder/
~~~ Then, run DART-ID in the python programming environment using the following command, specifying the path to the configuration file, example:

~~~
 dart_id -c path/to/config/example_config_file.yaml
~~~

### Evaluating quality of single cell preparation Timing: 1+ days

The quality of sample preparation should be evaluated for every single cell. The inclusion of negative control wells allows background signal to be characterized. Below we describe data analysis that can be implemented using the SCoPE2 GitHub[51] or Zendo[52] repositories. This SCoPE2 code was used to develop the scp Bioconductor package[53, 54], which can also be used to analyze the data.

68. **Filtering for quality peptide and protein identifications:** (? Troubleshooting)
  - Remove peptides with false discovery rate (FDR) *>* 1%. Sometimes, it may be desirable to filter out peptides with posterior error probabilities (PEP) *>* 0.02
  - Remove peptides belonging to proteins with FDR *>* 1%
  - Remove reverse and contaminant peptides (as annotated by MaxQuant)
  - Remove peptides with precursor intensity fraction (PIF) *<* 0.8
  - Remove peptides with *>* 10% reporter ion intensity of the carrier channel reporter ion intensity
69. **Data transformations:**
  - To control for differences in the sampling of elution profiles in different LC-MS/MS runs, a common reference was used as a denominator for all single-cell quantification. This means that the reporter ion intensity for every peptide in every single cell or control was divided by the reporter ion intensity from the reference channel from the same peptide.
  - Then, to control for differences in sample loading, the relative peptide reporter ion intensities (relative to reference) for each single cell or control were divided by their median reporter ion intensity.
  - Then, to put all peptides on the same scale, the relative peptide reporter intensities for each peptide across all single cells and controls were divided by their median value.
  - Protein quantification was determined by taking the median peptide value for peptides belonging to said protein (denoted by MaxQuant as Leading Razor Protein).
70 **Expected signal level:** The preparation of single cells can be evaluated by looking for the expected amount of ions observed. The expectation is that well-prepared cells yield reporter ion intensities roughly proportional with the intensity from the carrier or reference channels (so ratios of 200:1 or 5:1, respectively) and that control wells and poorly-prepared single cells yield values less than that ratio (200:0.1 or 5:0.1, for example).
71 **Consistency of quantification:** Additionally, the preparation of single cells can be evaluated by looking for the consistency of peptide quantification within those peptides coming from the same proteins. This is captured by the coefficient of variation statistic (CV), the expectation being that well-prepared cells yield peptides with consistent quantification if they come from the same proteins (typical interquartile range of protein CVs for a successful single cell would 0.2 to 0.4), and thus lower CV than the control wells or poorly-prepared single cells.
72 **Identifying failed wells:** For every single cell and control, plot the median relative reporter ion intensity versus the median CV. Failed single cells and negative controls should cluster apart from successfully-prepared single cells.
73 **Identifying batch effects:** As in any experiment, multiple parts of SCoPE2 are subject to potential batch effects. Experiments should be designed to make sure experimental conditions do not correlate with potential batch effects. For SCoPE2, potential batch effects include:
  - 384-well plate (i.e. don’t process different conditions on separate 384-well plates)
  - Plate location (i.e. edge effects)
  - Chromatography (i.e. LC-MS/MS column, or run to run variability)
  - TMT label (i.e. differences in labeling efficiency due to contaminated reagent)

## Data availability recommendations

- **Facilitating data reuse:** In addition to following conventional practices for deposition of raw mass spectrometry data and search results, we make the following recommendations for data availability:

Prepare data files from intermediate steps of data processing. We usually provide at least 3 files in comma separated values (csv) format as follows:

1. `Peptides-raw.csv –` peptides × single cells at 1 % FDR and including peptides identified by DART-ID. The first 2 columns list the corresponding protein identifiers and peptide sequences and each subsequent column corresponds to a single cell.
2. `Proteins-processed.csv –` proteins × single cells at 1 % FDR, imputed (K-nearest neighbors, k = 3) and batch corrected (ComBat package in R).
3. `Cells.csv –` annotation × single cells. Each column corresponds to a single cell and the rows include relevant metadata, such as, cell type if known, measurements from the isolation of the cell (e.g. from index sorting by FACS), and derivative quantities, i.e., rRI, CVs, reliability.

These files can be included with the raw file deposition, and as supplementary data provided with a manuscript.

### Troubleshooting

| Issue | Cause | Resolution | Step |
| --- | --- | --- | --- |
| **LC-MS system** |  |  |  |
| No signal from a SCoPE sample injection | Autosampler needle may not have aspirated fully the expected $1\mu\text{l}$ injection, resulting in an injection of air | Spin down sample and ensure that liquid is evenly settled at bottom of tube and/or service the autosampler. | (65) |
| Low number of identified peptides | Poor sample delivery, lower-than-expected carrier, contamination, suboptimal parameters or another problem. | This can be caused by many different factors. Look at the DO-MS report (do-ms.slavovlab.net) to specifically diagnose the problem. | (68) |
| Apex offset distribution not centered around 0 | Consistently sampling peptides before or after their elution apices | Adjust the length of your duty cycle by changing the number of peptides submitted for MS2 (top N setting) and/or the MS2 fill times. See Specht et al.<sup>25</sup> for examples and guidelines. | (21) |
| Increased pressure trace | Undigested protein or nucleic acids obstructing column | If using the recommended ionOpticks column, remove column, examine the end for obstructions under a light microscope, and trim end just beyond obstructions using a diamond cutter. | (21) |
| Multiple MS2 scans / peptide | Broad elution peak | Aging LC column or free space in the LC plumbing. | (21) |
| **Sample** |  |  |  |
| Poor labeling efficiency | TMT levels may be hydrolyzed or conditions may not be optimal for TMT labeling | Check your TMT stock and ensure the post-digest pH is approximately 8. | (39) |
| Relative Reporter Ion Ratios do not match expected values | Sample input may not match expected input amount or TMT labels may not have been properly quenched | Use the positive controls to diagnose the problem. | (68) |
| Low correlations between cell-type-based reporter ion intensity ratios for single-cell channels and carrier channels (if using an experimental design that features two carrier channels) | Possible cross-labeling of samples | Ensure that labeling and quenching conditions match those described here and that hydroxylamine is still active. Additionally, ensure that the isobaric carrier is well labeled. Unlabeled carrier peptides may be labeled by residual labels used for the single-cell channels if the labels have not been fully quenched. | (68) |
| **Analysis** |  |  |  |
| Poor labeling Efficiency | Suboptimal labeling conditions or hydrolyzed TMT | Check your TMT stock and ensure the post-digest pH is approximately 8. | (39) |

The authors delivered a workshop covering SCoPE2 experimental design, sample preparation and data analysis at the SCP2019 conference on single-cell proteomics, held in Boston in June 2019. Videos from this workshop are available on YouTube and may be of use to those seeking further clarification or troubleshooting^55^.

### Timings

#### Anticipated Results

The SCoPE2 method was initially applied to the study of U937 monocyte cells differentiating to macrophage-like cells, which provided an example of how the quantified proteins can be used for biological analysis^18^. Here, we exemplify the described SCoPE2 protocol (using cellenONE for cell isolation) with a smaller and simpler experiment (analyzing 6 SCoPE2 sets consisting of single HeLa and U937 cells) so that we can focus on key technical benchmarks that will be helpful for new groups adopting the SCoPE2 workflow.

The success and the problems of data acquisition can be analyzed by the corresponding DO-MS report. The full report consisting of many dozens of plots can be found as supplemental material, and Fig. 2 shows a few plots demonstrating expected results. The distribution of precursor intensities in Fig. 2a are useful benchmark to evaluate problems with sample pick up (which manifests as sporadic runs with very low precursor intensities) or poor sample delivery to the instrument, which manifests with decline in the precursor abundances relative to a reference standard. To make this comparison meaningful, DO-MS compares distributions composed of the same peptides across all runs^25^.

**Figure 2.**
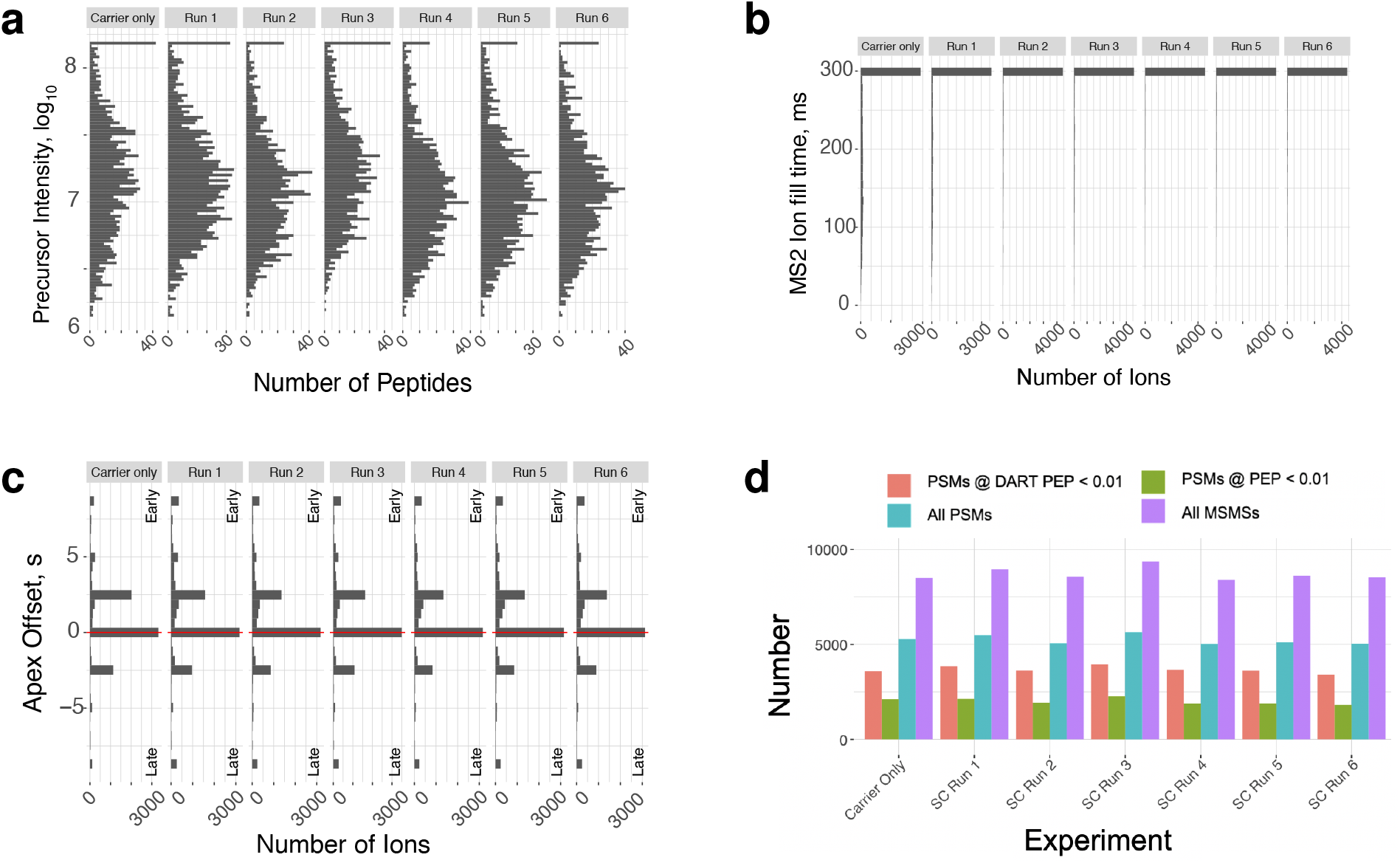
Evaluating data acquisition and interpretation using diagnostic plot generated by DO-MS. (**a**) Distributions of intensities for precursors quantified in all displayed experiments. (**b**) The distributions of times for accumulating ions for MS2 scans indicate that most ions were accumulated for maximum time allowed, 300ms. (**c**) Distributions of times between the apexes of elution peaks and the time when they were sampled for MS2 scans. (**d**) Number of MS2 scans and peptide spectrum matches (PSMs) at different levels of confidence, with and without DART-ID. These and many other plots are automatically generated by DOMS^25^.

Suboptimal choice of instrument parameters might result in short MS2 accumulation times and thus in sampling only a few copies from the single-cell peptides^19,22,25^. To evaluate and control this possibility, DO-MS displays the distributions of MS2 accumulation times as shown in Fig. 2b. As expected for our 6 runs, the isobaric carrier did not limit MS2 accumulation times, and peptides selected for MS2 scans were accumulated for the maximum time specified in our instrument methods, 300ms.

To maximize the copy number of ions sampled for quantification and the purity of MS2 spectra, SCoPE2 aims to sample elution peaks close to the apex^18^. To help achieve this goal, DO-MS displays the distributions of apex offsets as shown in Fig. 2c. In the 6 example runs included here, most ions were sampled within a few seconds from the apex without systematic bias of early or late sampling. If such biased are present, follow the recommendations outlined in ref.^25^ to mitigate them.

DO-MS evaluates the number of identified peptides in the context of total number of MS2 scans taken, the total number of peptide-spectrum-matched (PSMs), in addition to the number of confident PSMs based on spectra alone and based on both spectra and DART-ID. The example plot in Fig. 2 shows that most runs acquired about 9,000 MS2 scans (which is close to the number expected for full duty cycles) and a large fraction of these MS2 spectra were assigned sequences with high confidence, about 20 % based on spectra alone and about 40 % based on spectra and retention time.

Before evaluating the accuracy of protein quantification by SCoPE2, we evaluate whether the relative reporter ion (RI) intensities follow the expected trends. In particular, negative control wells shown have low RI intensities (low background) and the single-cell RI intensities should be about 1/x the RI intensities of the isobaric carrier, where x is the number of cells in the isobaric carrier. These expected results are illustrated in Fig. 3a with data from 5 single HeLa cells and 5 single U937 cells. The RI intensities for the HeLa cells are larger because of their larger cell sizes. Most RIs are not detected in the negative control, indicating that background noise is below the limit of detection of our system.

**Figure 3.**
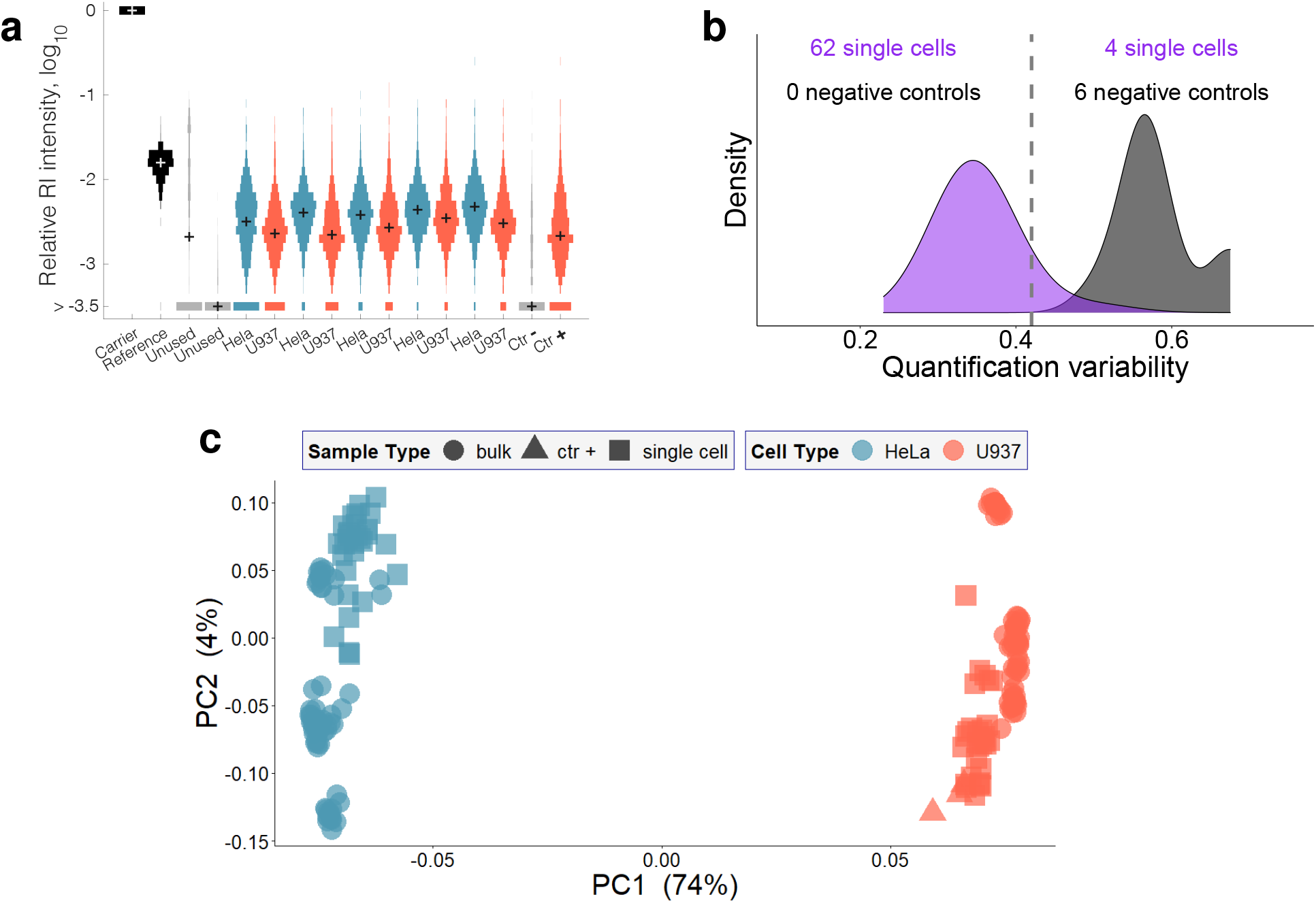
Evaluating protein quantification results from SCoPE2 analysis. (**a**) Distributions of relative reporter ion (RI) intensities for all samples from a single SCoPE2 set. The samples include carrier, reference, 5 single HeLa cells, 5 single U937 cells, a negative control well (Ctr −), and a positive control well (Ctr +) from diluted U937 cell lysate. Two of the TMT labels (between the reference and the single cells) are not used because they are affected by isotopic impurities of the TMT labels used for the carrier and the reference. The bin marked by *>* − 3.5 corresponds mostly to RI intensities below the detection limit (missing values). (**b**) Distributions of coefficients of variation (CVs) for the relative quantification of proteins based on different peptides originating from the same protein. The CVs are computed both for the single cells and the negative control wells. (**c**) Principal component analysis (PCA) of 126 single cells and bulk samples. The PCA is based on 1756 proteins with about 1,000 proteins quantified per single cell. Panels b and c are generated by the SCoPE2 pipeline^52^.

In a large scale experiment, some single cell samples may not provide useful data because of problems during cell isolation or sample preparation. Such samples can be identified based on the low consistency of quantification derived from different peptides as shown in Fig. 3b. This diagnostic plot is generated by the SCoPE2 analysis pipeline^18,52^.

Low dimensional projections, such as the principal component analysis (PCA) shown in Fig. 3b, are frequently used to summarize single-cell data, and the degree to which similar samples cluster together may reflect the discriminatory power of the data. However, samples may be separated by PCA because of technical signals, such as batch effects or incomplete normalization of the size difference between HeLa and U937 cells, Fig. 3a. To evaluate whether the separation along the first principal component (PC1) is indeed capturing cell-type specific differences in protein abundance, we strongly recommend including bulk samples, as in the PCA displayed in Fig. 3c. We expect single cells and bulk samples from the same cell type to cluster close to each other and far away from clusters corresponding to different cell types, as observed in Fig. 3c.

## Data availability

All data are available at MassIVE MSV000087041 and via the SCoPE2 website scope2.slavovlab.net/mass-spec/protocol

## Code availability

The SCoPE2 pipeline used here is available at github.com/SlavovLab/SCoPE2 and via the SCoPE2 website scope2.slavovlab.net/mass-spec/protocol

## Acknowledgments

We thank A. Chen, and J. Neveu for assistance, discussions and constructive comments. This work was funded by a New Innovator Award from the NIGMS from the National Institutes of Health to N.S. under Award Number DP2GM123497, an Allen Distinguished Investigator award through The Paul G. Allen Frontiers Group to N.S., a Seed Networks Award from CZI CZF2019-002424 to N.S., through a Merck Exploratory Science Center Fellowship, Merck Sharpe & Dohme Corp. to N.S., and a Thermo Scientific Tandem Mass Tag Research Award to

E.E. Funding bodies had no role in data collection, analysis, and interpretation.

## Author Contributions

**Experimental design**: A.P. and N.S.

**Sorting cells**: A.L. and A.P

**LC-MS/MS**: G.H, E.E., H.S. and D.H.P.

**Sample preparation**: A.P.

**Funding**: N.S, E.E.

**Data analysis**: A.P. and N.S.

**Supervision**: N.S.

**Writing & editing**: E.E., A.P. and N.S.

---

**Extended Data Figure 1.**
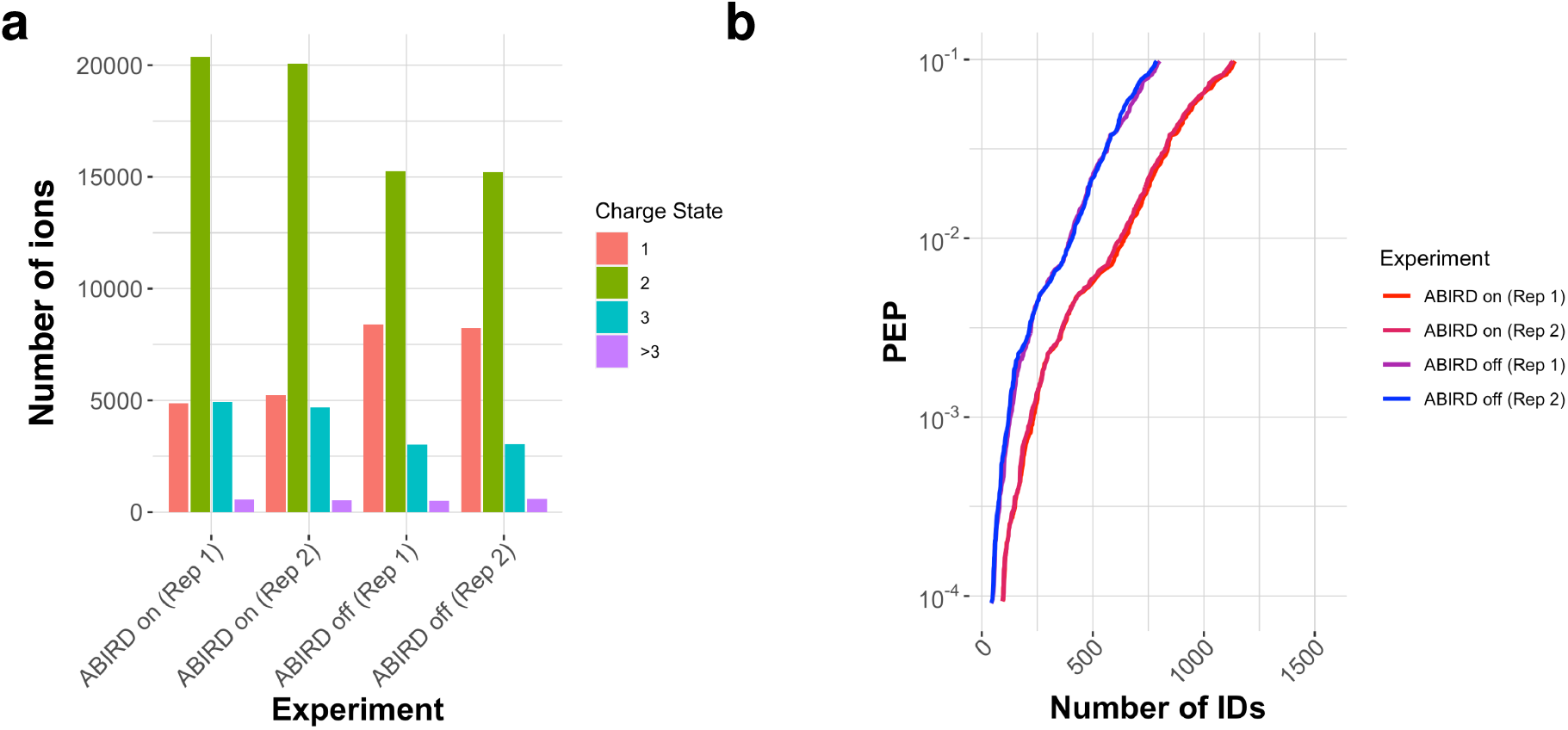
The ABIRD device may suppress contaminant ions and enhance peptide identification. ABIRD may suppress contaminant ions and enhance peptide identification. Replicate injections of an 1x Standard were analyzed with the ABIRD on or off. (**a**) The replicates with ABIRD on had a reduced number of +1 ions (likely corresponding to contaminants) and an increased number of higher charge state ions, which are likely to correspond to peptides. (**b**) With the ABIRD on, the number of identified peptides is increased across all confidence (PEP; posterior error probabilities).

## References

1. Symmons, O. & Raj, A. What’s Luck Got to Do with It: Single Cells, Multiple Fates, and Biological Nondeterminism. Molecular cell 62, 788–802 (2016).

2. Paul, I., White, C., Turcinovic, I. & Emili, A. Imaging the future: the emerging era of single-cell spatial proteomics. en. FEBS J. (Dec. 2020).

3. Levy, E. & Slavov, N. Single cell protein analysis for systems biology. Essays In Biochemistry 62. doi:10.1042/EBC20180014 (4 2018).

4. Regev, A. et al. Science forum: the human cell atlas. Elife 6, e27041 (2017).

5. Ziegenhain, C. et al.. Comparative analysis of single-cell RNA sequencing methods. Molecular cell 65, 631–643 (2017).

6. Engblom, C. et al.. Osteoblasts remotely supply lung tumors with cancer-promoting SiglecFhigh neutrophils. en. Science 358 (Dec. 2017).

7. Shaffer, S. M. et al.. Rare cell variability and drug-induced reprogramming as a mode of cancer drug resistance. Nature 546, 431–435 (2017).

8. Franks, A., Airoldi, E. & Slavov, N. Post-transcriptional regulation across human tissues. PLoS computational biology 13, e1005535 (2017).

9. Slavov, N. Unpicking the proteome in single cells. Science 367, 512–513. ISSN: 0036-8075 (2020).

10. Washburn, M. P., Wolters, D. & Yates, J. R. Large-scale analysis of the yeast proteome by multidimensional protein identification technology. Nature biotechnology 19, 242–247 (2001).

11. Altelaar, A. F. M., Munoz, J. & Heck, A. J. R. Next-generation proteomics: towards an integrative view of proteome dynamics. en. Nat. Rev. Genet. 14, 35–48 (Jan. 2013).

12. Cravatt, B. F., Simon, G. M. & Yates Iii, J. R. The biological impact of mass-spectrometry-based proteomics. Nature 450, 991 (2007).

13. Boersema, P. J., Raijmakers, R., Lemeer, S., Mohammed, S. & Heck, A. J. Multiplex peptide stable isotope dimethyl labeling for quantitative proteomics. Nature protocols 4, 484–494 (2009).

14. Muntel, J. et al.. Comparison of Protein Quantification in a Complex Background by DIA and TMT Workflows with Fixed Instrument Time. Journal of Proteome Research 18. PMID: 30726097, 1340–1351 (2019).

15. Slavov, N. Single-cell protein analysis by mass spectrometry. Current Opinion in Chemical Biology 60, 1–9. ISSN: 1367-5931 (2020).

16. Kelly, R. T. Single-Cell Proteomics: Progress and Prospects. Molecular & Cellular Proteomics 19, 1739–1748 (2020).

17. Budnik, B., Levy, E., Harmange, G. & Slavov, N. SCoPE-MS: mass-spectrometry of single mammalian cells quantifies proteome heterogeneity during cell differentiation. Genome Biology 19, 161 (2018).

18. Specht, H. et al.. Single-cell proteomic and transcriptomic analysis of macrophage heterogeneity using SCoPE2. Genome Biology 22. doi:10.1186/s13059-021-02267-5 (2021).

19. Specht, H. & Slavov, N. Optimizing Accuracy and Depth of Protein Quantification in Experiments Using Isobaric Carriers. Journal of Proteome Research 20. PMID: 33190502, 880– 887 (2021).

20. Specht, H. et al.. Automated sample preparation for high-throughput single-cell proteomics. bioRxiv 10.1101/399774. doi:10.1101/399774. https://doi.org/10.1101/399774 (2018).

21. Leduc, A., Huffman, R. G. & Slavov, N. Droplet sample preparation for single-cell proteomics applied to the cell cycle. bioRxiv 2021.04.24.441211. doi:10.1101/2021.04.24.441211 (2021).

22. Cheung, T. K. et al.. Defining the carrier proteome limit for single-cell proteomics. Nature Methods 18, 76–83 (2021).

23. Marx, V. A dream of single-cell proteomics. Nature Methods 16, 809–812. ISSN: 1548-7105 (2019).

24. Chen, A. T., Franks, A. & Slavov, N. DART-ID increases single-cell proteome coverage. PLOS Computational Biology 15, 1–30 (July 2019).

25. Huffman, G., Chen, A. T., Specht, H. & Slavov, N. DO-MS: Data-Driven Optimization of Mass Spectrometry Methods. J. of Proteome Res. doi:10.1021/acs.jproteome.9b00039 (2019).

26. Bendall, S. C. et al.. Single-cell mass cytometry of differential immune and drug responses across a human hematopoietic continuum. Science 332, 687–696 (2011).

27. Albayrak, C. et al.. Digital quantification of proteins and mRNA in single mammalian cells. Molecular cell 61, 914–924 (2016).

28. Darmanis, S. et al.. Simultaneous multiplexed measurement of RNA and proteins in single cells. Cell reports 14, 380–389 (2016).

29. Hughes, A. J. et al.. Single-cell western blotting. Nature methods 11, 749–755 (2014).

30. Specht, H., Emmott, E., Perlman, D. H., Koller, A. & Slavov, N. High-throughput single-cell proteomics quantifies the emergence of macrophage heterogeneity. bioRxiv. doi:10.1101/665307 (2019).

31. Virant-Klun, I., Leicht, S., Hughes, C. & Krijgsveld, J. Identification of maturation-specific proteins by single-cell proteomics of human oocytes. Molecular & Cellular Proteomics 15, 2616–2627 (2016).

32. Lombard-Banek, C., Moody, S. A. & Nemes, P. Single-Cell Mass Spectrometry for Discovery Proteomics: Quantifying Translational Cell Heterogeneity in the 16-Cell Frog (Xenopus) Embryo. Angewandte Chemie International Edition 55, 2454–2458 (2016).

33. Cong, Y. et al.. Improved Single-Cell Proteome Coverage Using Narrow-Bore Packed NanoLC Columns and Ultrasensitive Mass Spectrometry. Analytical Chemistry 92, 2665–2671 (Feb. 2020).

34. Cong, Y. et al.. Ultrasensitive single-cell proteomics workflow identifies > 1000 protein groups per mammalian cell. Chemical Science (2020).

35. Brunner, A.-D. et al.. Ultra-high sensitivity mass spectrometry quantifies single-cell proteome changes upon perturbation. bioRxiv (2020).

36. Zhu, Y. et al.. Nanodroplet processing platform for deep and quantitative proteome profiling of 10–100 mammalian cells. Nature communications 9, 882 (2018).

37. Dou, M. et al.. High-throughput single cell proteomics enabled by multiplex isobaric labeling in a nanodroplet sample preparation platform. Analytical chemistry 91, 13119–13127 (2019).

38. Li, Z.-Y. et al.. Nanoliter-scale oil-air-droplet chip-based single cell proteomic analysis. Analytical chemistry 90, 5430–5438 (2018).

39. Liang, Y. et al.. Fully Automated Sample Processing and Analysis Workflow for Low-Input Proteome Profiling. en. Anal. Chem. 93, 1658–1666 (Jan. 2021).

40. Lafzi, A., Moutinho, C., Picelli, S. & Heyn, H. Tutorial: guidelines for the experimental design of single-cell RNA sequencing studies. Nature Protocols 13, 2742–2757 (2018).

41. Barkas, N. et al.. Joint analysis of heterogeneous single-cell RNA-seq dataset collections. Nature methods 16, 695–698 (2019).

42. Marx, V. A dream of single-cell proteomics. Nature Methods. doi:10.1038/s41592-019-0540-6 (Dec. 2019).

43. Grün, D. & Van Oudenaarden, A. Design and Analysis of Single-Cell Sequencing Experiments. Cell 163, 799–810 (2015).

44. Meier, F. et al.. Online Parallel Accumulation-Serial Fragmentation (PASEF) with a Novel Trapped Ion Mobility Mass Spectrometer. Molecular & Cellular Proteomics 17, 2534–2545 (Jan. 2018).

45. Specht, H. & Slavov, N. Transformative opportunities for single-cell proteomics. Journal of Proteome Research 17, 2563–2916 (8 June 2018).

46. Cox, J. & Mann, M. MaxQuant enables high peptide identification rates, individualized ppb-range mass accuracies and proteome-wide protein quantification. Nature biotechnology 26, 1367–1372 (2008).

47. Tyanova, S., Temu, T. & Cox, J. The MaxQuant computational platform for mass spectrometry-based shotgun proteomics. Nature protocols 11, 2301 (2016).

48. Yu, S.-H., Kyriakidou, P. & Cox, J. Isobaric matching between runs and novel PSM-level normalization in MaxQuant strongly improve reporter ion-based quantification. Journal of proteome research 19, 3945–3954 (2020).

49. Fondrie, W. E. & Noble, W. S. mokapot: Fast and Flexible Semisupervised Learning for Peptide Detection. Journal of Proteome Research 0, null (0).

50. Milo, R., Jorgensen, P., Moran, U., Weber, G. & Springer, M. BioNumbers-the database of key numbers in molecular and cell biology. Nucleic acids research 38, D750–D753 (2010).

51. Specht, H. et al.. Single-cell proteomic and transcriptomic analysis of macrophage hetero-geneity. GitHub, github.com/SlavovLab/SCoPE2 (2019).

52. Specht, H. et al.. Single-cell proteomic and transcriptomic analysis of macrophage heterogeneity using SCoPE2. Zenodo, 10.5281/zenodo.4339954 (2020).

53. Vanderaa, C. & Gatto, L. Utilizing Scp for the analysis and replication of single-cell proteomics data. bioRxiv. doi:10.1101/2021.04.12.439408 (2021).

54. Vanderaa, C. & Gatto, L. Mass Spectrometry-Based Single-Cell Proteomics Data Analysis. Bioconductor, 10.18129/B9.bioc.scp (2020).

55. SCP2019 Workshop on single-cell proteomics Aug. 2019. http://workshop2019.single-cell.net.

